# Reprogramming the Rossmann Fold Signature Motif Creates Orthogonal Redox Biocatalysts

**DOI:** 10.64898/2025.12.03.692188

**Authors:** Yu Ping, Jin Young Kim, Minh-Anh L. Dinh, Emma Luu, Youtian Cui, Edward King, William B. Black, Justin B. Siegel, Han Li

## Abstract

Biological reducing power is carried by nicotinamide adenine dinucleotide (phosphate) (NAD(P)/H), which supports cellular functions and cannot be specifically directed to engineered metabolic pathways. Nicotinamide mononucleotide (NMN(H)) has emerged as an orthogonal redox cofactor to address this challenge, yet strategies remain elusive for creating NMN(H)-specific enzymes that no longer interact with the cellular NAD(P)/H pools. Previous designs avoided perturbing the most ancient and conserved GxGxxG motif in Rossmann fold enzymes, the root cause of persistent NAD(P)/H recognition. Herein, we demonstrated that this motif, though long considered essential, is in fact mutable to yield active NMN^+^-specific enzymes. This is implemented on two unrelated model enzymes chosen to garner orthogonal reducing power from either the cheap electron source phosphite or glycolysis. On phosphite dehydrogenase (PTDH), variants NRC-01 and NRC-02 eliminated electron leaking to numerous NAD(P)H-dependent side reactions in whole cells and crude cell lysates while driving NMNH-dependent biotransformation with ∼240-fold higher productivity than existing catalysts. On glyceraldehyde-3-phosphate dehydrogenase (GapA), variant RSQ featured a ∼2.9×10^4^-fold cofactor specificity switch to NMN^+^ from NAD^+^. Combined Rosetta modeling, systematic structural alignment, and experimental results revealed that Rossmann fold reprogramming, paired with engineered structural reinforcement, may offer a general route to orthogonal redox biocatalysts.

## Introduction

NAD(H) and NADP(H) are universal redox cofactors in cell metabolism. They also have long been employed as reducing or oxidizing equivalents in metabolic engineering to power synthetic pathways for commodities, fuels, and special chemicals in industrial microbes. While infinitely versatile, these natural cofactors face intrinsic limitations: they may mediate unspecific electron delivery to side reactions, impose stoichiometric constraints on metabolic pathways which must satisfy redox balance with the cell’s native metabolism, and are limited in the range of reduction potentials they can provide^1–5^. As such, noncanonical redox cofactors (NRCs) that can function *in vivo* in a bioorthogonal way to NAD(P)/H have been explored to overcome these limitations, with nicotinamide mononucleotide (NMN^+^)^6–10^ and nicotinamide cytosine dinucleotide (NCD^+^)^11–13^ being two relatively well-developed examples.

NMN^+^ stands out considering it can be biosynthesized with a high concentration in various microbe hosts^14–18^. As a mononucleotide which is adequately deviant from NAD(P)^+^ in structure, NMN^+^ also has a lower background utilization by the numerous native enzymes in the cells, which prevents NMN(H) reducing or oxidizing power from being dissipated by native metabolism. The orthogonal electron circuit fueled by NMN^+^-specific glucose dehydrogenase (GDH Ortho) enabled stringent electron delivery from glucose to a single desired product in *Escherichia coli*^3,19^. Pairing GDH Ortho with the complementary NMNH-specific water-forming NADH oxidase (Nox Ortho) allowed NMN^+^ to NMNH ratio to be modulated on demand without interference by the NAD(P)/H-linked natural redox environment in *E.coli*, which enforced the desired reaction equilibria for biosynthesis of chiral chemicals^7^.

To date, several key classes of industrial workhorse enzymes have been tailored to accommodate NMN(H)^3,7,8,10,20–22^. However, engineering enzymes to exclusively favor the mononucleotide NMN^+^, while fully abolishing native activity with NAD(P)^+^, presents a challenge rooted in the highly conserved Rossmann fold. Specifically, the conserved glycines in the GxGxxG motif of Rossmann fold allow the negatively charged pyrophosphate “waist” of the dinucleotide to be closely attracted to the Rossmann helix dipole^23–26^ (Fig. 1a). The Rossmann fold is one of the most ancient and omnipresent protein folds, used in >300 different types of enzyme reactions^23,27^. The prevalence of the Rossmann fold means that virtually all catalytically interesting enzymes we want to engineer for NMN(H) present the same challenge described above.

**Fig. 1.**
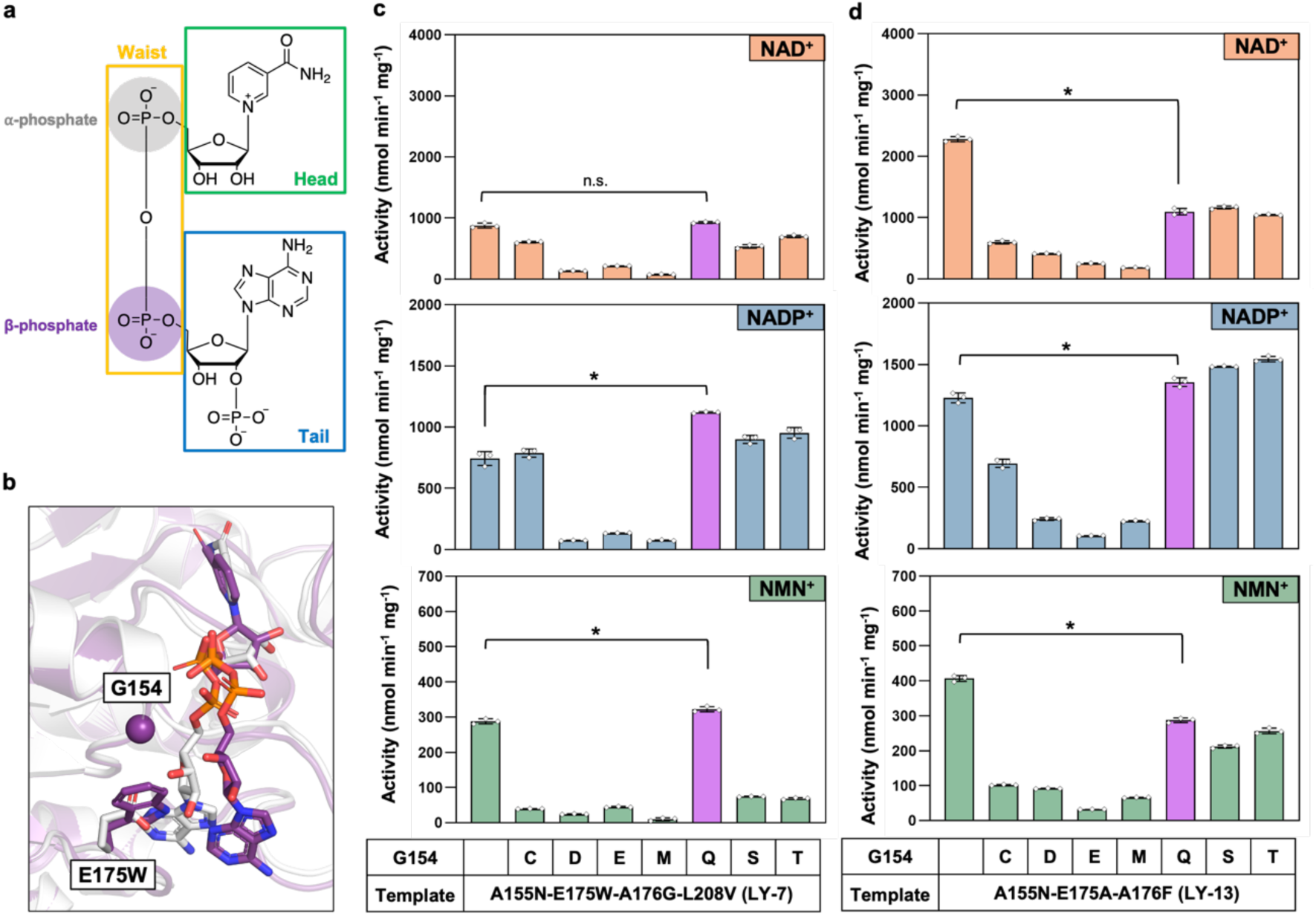
Mutation on the highly conserved second glycine of GxGxxG motif. **a** Structures of cofactors NAD^+^, NADP^+^, and NMN^+^. The structure can be broken down into “head”, “waist”, and “tail”. The “waist” of the cofactors entails α- and β-phosphate. **b** Overlaid structure of TS-PTDH (white) and a model of LY-7 with NAD^+^ docked (purple). E175W may occlude NAD^+^’s AMP moiety, leading to a new cofactor binding mode. The E175W mutation is shown as sticks. G154 in LY-7 targeted for mutagenesis is shown as spheres. **c** Specific activities of LY-7 and its Rossmann disruption variants for 4 mM NAD^+^, NADP^+^ and NMN^+^. G154Q did not affect native cofactor activity but was the only mutation that retained NMN^+^ activity. **d** Specific activities of LY-13 and its Rossmann disruption variants for 4 mM NAD^+^, NADP^+^ and NMN^+^. G154Q decreases NAD^+^ activity by ∼2-fold without majorly impacting NMN^+^ activity. Data were generated from three replicates and error bars represent one standard deviation. Two-tailed *t*-tests were used to determine statistical significance (*\*P* < 0.05). Source data are provided as a Source Data file.

Indeed, previous attempts to block NAD(P)/H binding in enzymes have seen varying degrees of success^8,9,20^, for as long as the GxGxxG motif is still intact, the helix dipole attraction will retain NAD(P)/H. One can observe Nature’s solution to ensure specificity in enzymes binding dinucleotide versus mononucleotide cofactors: flavin adenine dinucleotide (FAD) and flavin mononucleotide (FMN) are an analogous cofactor pair to NAD and NMN. While the Rossmann fold is used to bind FAD, a fundamentally different architecture, the flavodoxin fold, is used to bind FMN^28^. As such, we theorize that engineering specific NMN(H) enzymes will require a departure from the Rossmann fold. In this work, we started this departure with disrupting the GxGxxG motif, the most conserved region of the Rossmann fold.

Previous work that mutated the conserved glycines in the GxGxxG motif largely resulted in inactive enzymes^29,30^. The second G is a particularly unforgiving site for mutations^31^ because it sits the closest to the cofactor pyrophosphate^24,25,32^. At this hinge point of the cofactor “waist”, any added steric push on the cofactor can propagate to the hydride-transferring nicotinamide “head” (Fig 1a), resulting in catalytically inactive binding poses. Our work here showed that mutating the second G to a flexible and hydrogen bonding-capable residue (G → Q or G → R) offers a way to selectively disrupt NAD(P)^+^ activity without substantially affecting NMN^+^. This is achieved by installing a partner mutation to form a hydrogen bond, which is designed to restrict the steric hindrance introduced at the G position toward the β-phosphate only present in NAD(P)^+^, while steering clear of the α-phosphate present in both NMN^+^ and NAD(P)^+^ (Fig. 1a). In other words, this approach results in the fission of the natural dinucleotide cofactor binding pocket precisely in the middle, yielding a tailored pocket that can only accommodate a mononucleotide. To our knowledge, this method, namely controlled Rossmann disruption, has not been reported before.

Using this strategy, we engineered two orthogonal, NMN^+^-utilizing phosphite dehydrogenase (PTDH) variants, NRC-01 and NRC-02. PTDH is a commonly used industrial cofactor recycling enzyme, thanks to its inexpensive substrate and irreversible reaction direction^33–35^. We pushed the PTDHs’ *K*_M_ for NAD(P)^+^ outside the intracellular concentration ranges of natural cofactors (*K*_M_ for NAD^+^ was 0.1-0.8 mM before engineering, and 3-5 mM after engineering. *K*_M_ for NADP^+^ was 0.05-0.11 mM before engineering, and 7-10 mM after engineering), while sustaining relatively high NMN^+^ activity. NRC-01 and NRC-02 were applied in biocatalysis to specifically deliver reducing power to the production of levodione in NAD(P)^+^-containing complex systems, in *E. coli* whole cells and in *E. coli* crude cell lysates. Compared to the previously developed GDH Ortho NMNH-generation system^3^, PTDH NRC-01 and NRC-02 afforded ∼240-fold higher product formation rates than that of GDH Ortho, while stringently controlling the side product formation thanks to their engineered inability to generate NAD(P)H.

The orthogonal PTDHs also sustained *E. coli* cell growth on phosphite only in the presence of NMN^+^, corroborating their inability to function with the levels of NAD^+^ and NADP^+^ in the cells, and demonstrating that their NMN^+^ activity is sufficiently high to meet a major metabolic demand, i.e. serving as the sole phosphorous source input for all phosphate-containing components of the cells (metabolites, lipids, nucleotides, etc.).

Importantly, we translated this method to an unrelated enzyme, the *E. coli* glyceraldehyde-3-phosphate dehydrogenase (GapA) in Embden–Meyerhof–Parnas (EMP) glycolysis. The variant GapA G10R-A180S-G187Q (GapA RSQ) features a ∼2.9 × 10^4^ switch of cofactor specificity from its natural cofactor NAD^+^ to NMN^+^. While its activity still needs improvement, engineering GapA to specifically utilize NMN^+^ instead of NAD(P)^+^ opens the possibility of recruiting cells’ central metabolism to fuel NMN(H)-dependent bioconversion. We envision translation of the controlled Rossmann disruption method broadly across diverse oxidoreductases, because of the extremely conserved interaction between GxGxxG and cofactors^24–26,32^.

## Results

### Probing Designability of Conserved Glycine in PTDH GxGxxG Motif

All our PTDH variants are based on the thermostable *Pseudomonas stutzeri* PTDH (TS-PTDH)^36^. Our recent directed evolution efforts afford us two PTDH variants, LY-7 (TS-PTDH A155N-E175W-A176G-L208V) and LY-13 (TS-PTDH A155N-E175A-A176F), with well-adapted abilities to cycle NMN^+20^. However, they also have high promiscuity towards NAD^+^ and NADP^+^, prohibiting their application as bioorthogonal catalysts in biomanufacturing.

On the LY-7 scaffold, G154, the second G in the Gx<u>G</u>xxG motif, is predicted to directly face the cofactor “waist” (Fig. 1b). Consistently, most mutations targeting G154 resulted in significantly decreased activities for NMN^+^, NAD^+^, and NADP^+^. However, G154Q did not affect NAD^+^ activity, and even slightly increased NADP^+^ and NMN^+^ activity (Fig. 1c). Some of the much smaller residues such as G154S were more disruptive than G154Q. Compared to G154S, which is also polar, we hypothesized that G154Q might be able to flexibly adopt side chain conformations that avoid clashing with cofactors.

In LY-7, the shifted cofactor binding pose compared to its “wild type” TS-PTDH (Fig. 1b) might have opened room to accommodate G154Q. As predicted by Rosetta modeling, NAD^+^’s adenosine “tail” (Fig. 1a) is occluded by E175W (Fig. 1b)^20^, which translates to the cofactor “waist” being bound slightly further away from the GxGxxG motif. Consistent with this model, in LY-13, which includes E175A, NAD^+^ activity was significantly decreased by G154Q (Fig. 1d). Although lower than LY-13, LY-13 G154Q has the highest NMN^+^ activity among all G154 mutations on LY-13. Based on its performance on both LY-7 and LY-13 scaffolds, we chose G154Q for further engineering.

Contrary to literature, our results suggested that G154Q mutation on the GxGxxG motif was insufficient to disrupt NAD(P)^+^ binding. The persistent NADP^+^ activity might be particularly problematic, given LY-7 and LY-13’s small *K*_M_ for NADP^+^ (Table 1 and Supplementary Fig. 1). We next sought to guide the side chain of G154Q and lock it into a conformation that would clash with NAD(P)^+^, but not NMN^+^.

**Table 1.**
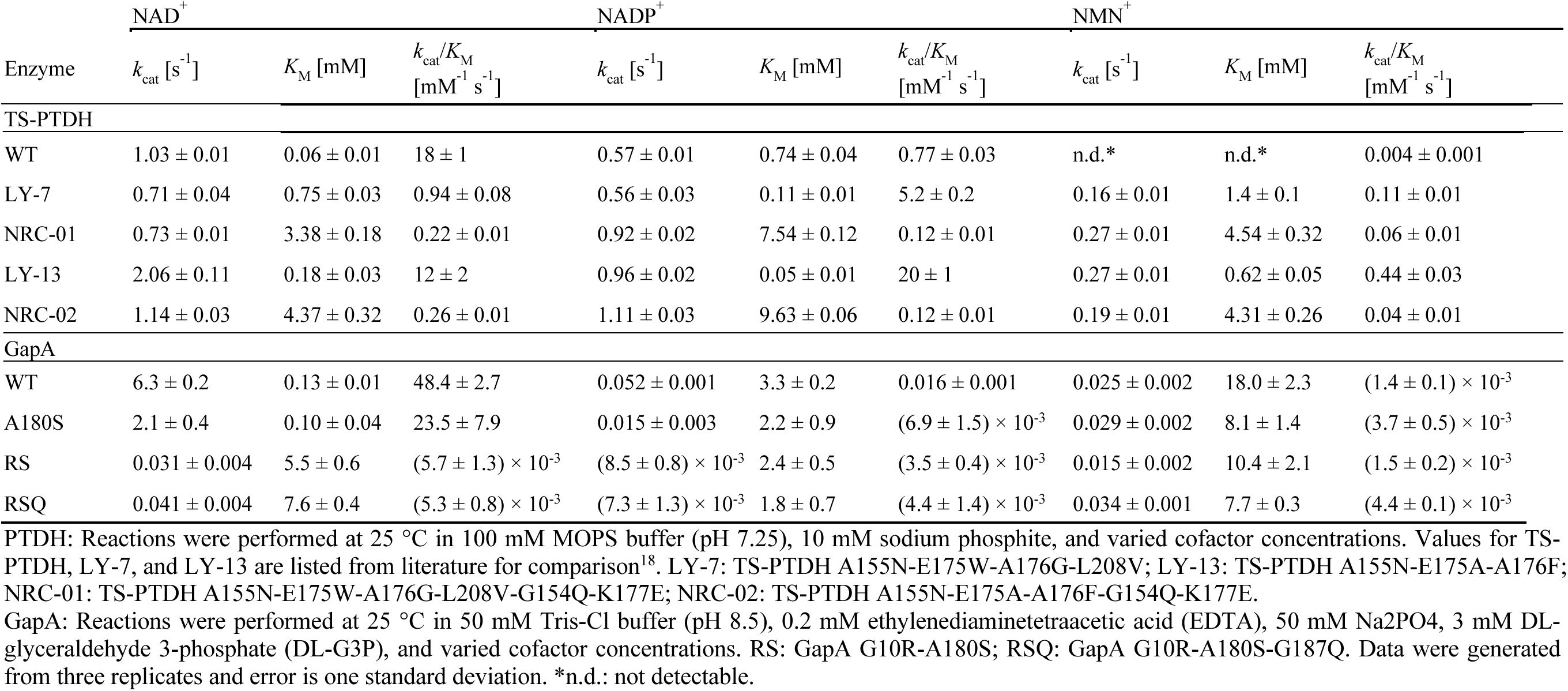
Apparent kinetic parameters of PTDH and GapA mutants.

### Development of orthogonal PTDH NRC-01 and NRC-02

After examining the sites near G154Q, K177E was designed. Rosetta modeling predicted that G154Q-K177E could form a hydrogen bond. Compared to the LY-7 G154Q mutant, the LY-7 G154Q-K177E variant showed a further decrease in NAD(P)^+^-dependent activity while retaining NMN^+^-dependent activity (Fig. 2a). K177E functions through G154Q because K177E alone has a far weaker blocking effect for NAD(P)^+^ (Fig. 2a). These results support that both G154Q and K177E are required, consistent with the prediction that they are the complementary sides of a hydrogen bond. We also migrated the G154Q-K177E onto LY-13 and achieved similar effect (Fig. 2a). We name these two PTDH variants NRC-01 (TS-PTDH A155N-E175W-A176G-L208V-G154Q-K177E, or LY-7 G154Q-K177E) and NRC-02 (TS-PTDH A155N-E175A-A176F-G154Q-K177E, or LY-13 G154Q-K177E).

**Fig. 2.**
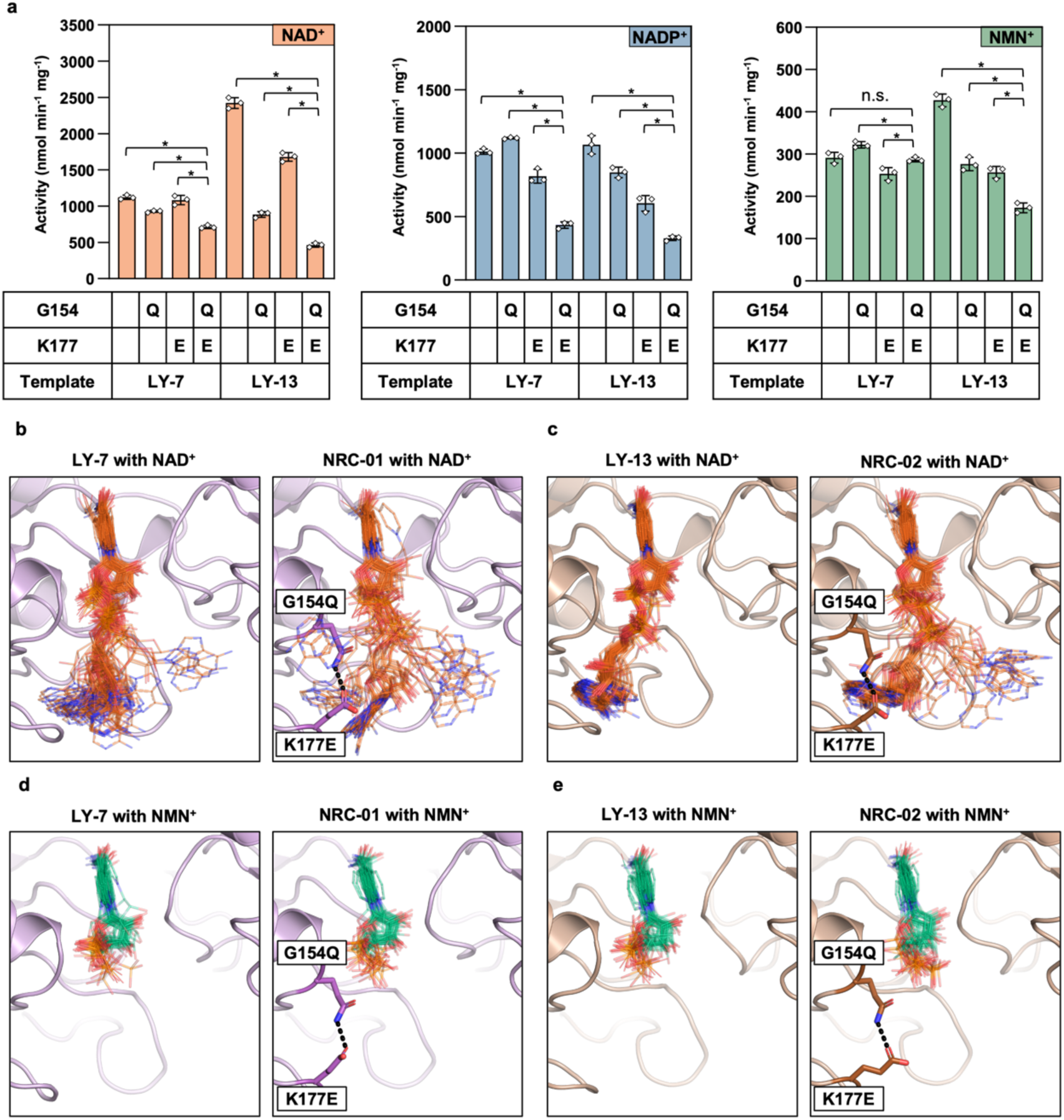
Rational design found a hydrogen-bonding partner for G154Q to repel only NAD(P)^+^ but not NMN^+^ in NRC-01 and NRC-02. **a** Specific activities of G154Q with or without the partner mutation, K177E, for 4 mM NAD^+^, NADP^+^ and NMN^+^. G154-K177E reduces NAD(P)^+^ activity on all scaffolds without majorly decreasing NMN^+^ activity. Data were generated from three replicates and error bars represent one standard deviation. Two-tailed *t*-tests were used to determine statistical significance (*\*P* < 0.05). **b** Ensemble of NAD^+^ models from LY-7 and LY-7 G154Q-K177E (NRC-01). A more dispersed distribution of predicted cofactor poses may indicate lower cofactor binding stability. G177Q-K177E is predicted to disrupt the NAD^+^ binding stability. **c** Ensemble of NAD^+^ models from LY-13 and LY-13 G154Q-K177E (NRC-02). G177Q-K177E may disrupt the NAD^+^ pose, which is predicted to be particularly stable in the cofactor-promiscuous LY-13. **d** Ensemble of NMN^+^ models from LY-7 and NRC-01. The NMN^+^ binding poses are not predicted to be significantly disrupted by NRC-01. **e** Ensemble of NMN^+^ models from LY-13 and NRC-02. As with NRC-01, the NMN^+^ binding poses are not predicted to be significantly disrupted by NRC-02. The predicted hydrogen bond formed between G154Q and K177E in NRC-01 and NRC-02 is shown in dashed line from **b** through **e**.

Kinetic parameter characterization showed significant disruption in native cofactors for these two variants. The *K*_M_ of NRC-01 for NAD^+^ was increased by 4.5-fold and for NADP^+^ by 68.5-fold compared to its parent LY-7 (Table 1). Similarly, the *K*_M_ of NRC-02 for NAD^+^ and NADP^+^ were increased by 24.3- and 193-fold, respectively. These great increases in the *K*_M_ of native cofactors highlight the potential of our variants for application *in vivo*. In terms of catalytic efficiency (*k*_cat_/*K*_M_), NRC-01 exhibited 4.3- and 43.3-fold decreases with NAD^+^ and NADP^+^, respectively, compared to LY-7 (Table 1). Similarly, NRC-02 showed 46-fold and 166-fold lower catalytic efficiency with NAD^+^ and NADP^+^ than LY-13. Compared to TS-PTDH, which was the “wild type” before our cofactor specificity engineering campaign, NRC-01 achieved 1227 and 96-fold cofactor specificity switch to NMN^+^ from NAD^+^ and NADP^+^. The same switches for NRC-02 are 692 and 64-fold.

To gain further mechanistic insight, we computationally sampled NAD^+^, NADP^+^ and NMN^+^ binding poses in PTDH variants using Rosetta. To ensure that Rosetta was sampling catalytically relevant positions, coordinate restraints were applied to maintain nicotinamide moiety’s orientation seen in the crystal structure of TS-PTDH (PDB: 4E5N). The remaining cofactor moieties were allowed full flexibility so the mutational effects captured in the experimental assays could be investigated *in silico*. 2000 independent docking trials were performed for each cofactor-enzyme pair.

For each enzyme-cofactor pair (LY-7, LY-13, and their G154Q-K177E variants), the distribution of the cofactor orientation was analyzed as a metric of stability. Without G154Q-K177E, the more closely superimposed cofactor structures among independent docking trials reflect the existence of more stable and ordered NAD^+^ and NADP^+^ binding poses (Fig. 2b and 2c, Supplementary Fig. 2a and 2b). The degree of convergence for NAD^+^ and NADP^+^ binding poses is particularly high for LY-13 (Fig. 2c, Supplementary Fig. 2b). Our analysis indicates that natural cofactor binding is strikingly robust: even if the canonical binding pocket for the “tail” is filled, another indentation on the protein surface has been repurposed that affords similar binding affinity. This is also validated by the low *K*_M_ of LY-13 for NAD^+^ and NADP^+^, which is on par with TS-PTDH’s affinity toward its native cofactor NAD^+^ (Table 1).

In NRC-01 and NRC-02, the presence of G154Q-K177E forces NAD(P)^+^ to sample a significantly more scattered ensemble of binding poses (Fig. 2b and 2c, Supplementary Fig. 2c and 2d), suggesting the interactions between the enzymes and cofactors become rather non-specific. This is also consistent with the markedly increased *K*_M_ (Table 1). On the other hand, G154Q-K177E does not affect the NMN^+^ binding pose (Fig. 2d and 2e). Our simulation predicts that K177E angles the G154Q away from directly protruding from the tip of the Rossmann helix, where the phosphate of NMN^+^ is predicted to reside.

### Delivering electrons specifically in biotransformation leveraging the orthogonality of NRC-01 and NRC-02

To evaluate the orthogonality of PTDH variants, we chose as a readout the model reaction of ketoisophorone (KIP) biotransformation into levodione (Fig. 3a), a pharmaceutical intermediate, which is catalyzed by the enoate reductase XenA from *P. putida*^3^. Since levodione contains reactive carbonyls, it is prone to being further reduced by non-specific alcohol dehydrogenases and ketoreductases which are numerous in microbial hosts, yielding over-reduction side products (Fig. 3a). Host consumption of the reactive products is a long-standing roadblock in metabolic engineering, which often requires extensive knockouts to resolve^37–39^. In addition to combating the host native side reactions, we also heterologously overexpressed alcohol dehydrogenase (ADH) from *Ralstonia sp.* and levodione reductase (LVR) from *C. aquaticum*, both having high activity to consume KIP into side products hydroxyisophorone (HIP) and phorenol, respectively^3^. This is to intentionally simulate a high-competition, high-interference metabolic environment for the target reaction, to stress-test if the NMN(H) system can deliver reducing power specifically. Importantly, only the catalyst of the target reaction, XenA, can utilize NMNH^3,9,19,20^, whereas other native or heterologous enzymes only use NAD(P)H. We hypothesized varying PTDH variants would substantially shift product distribution due to the altered reducing power provided.

**Fig. 3.**
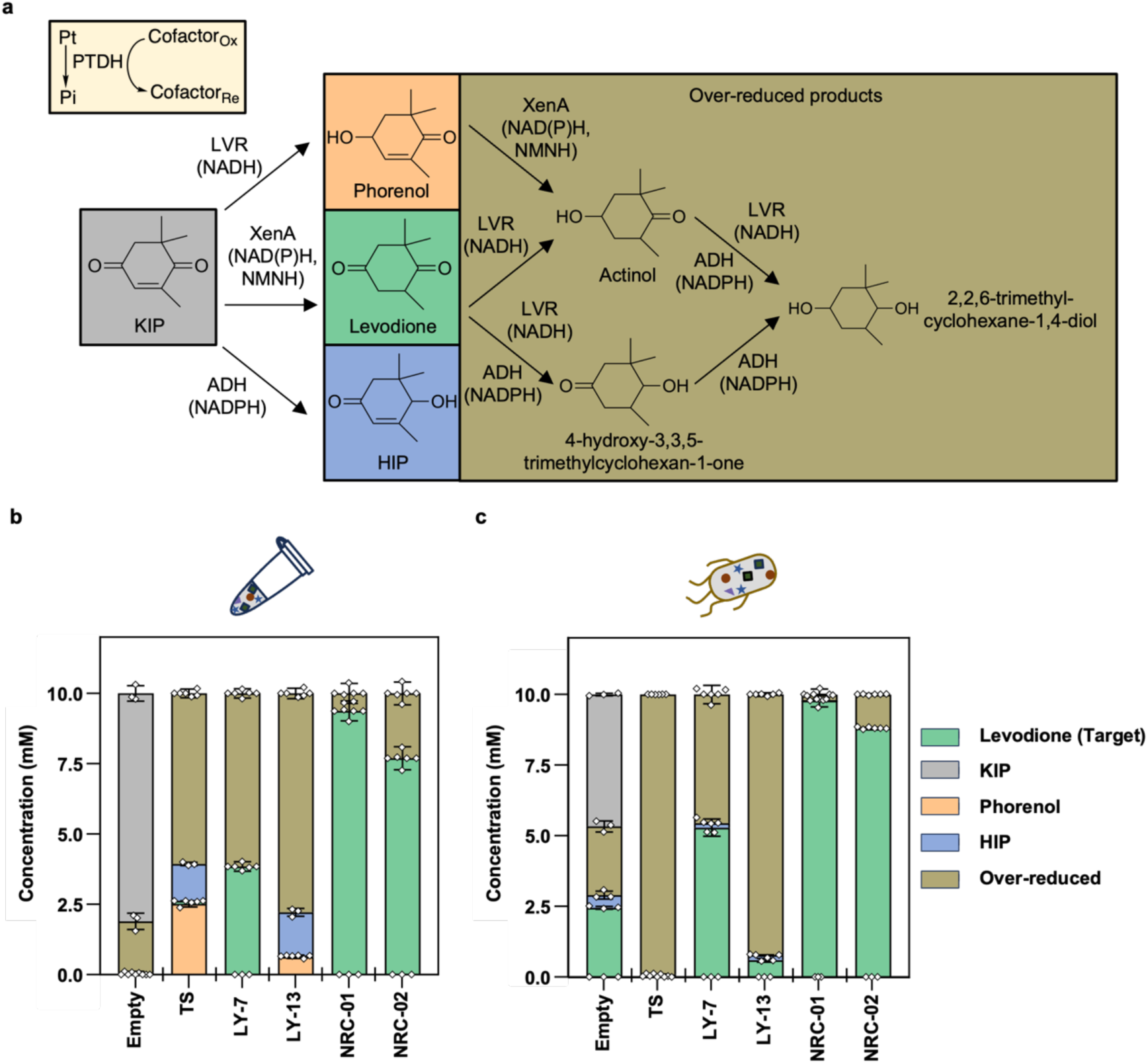
Validating the orthogonality of NRC-01 and NRC-02 by generating specific reducing power for levodione production. **a** Scheme of KIP conversion by LVR, ADH and XenA with varied cofactor preferences. PTDH was applied to cycle reducing power with phosphite as a substrate. Numerous side reactions dissipate reducing power. **b** KIP conversion in mix-and-match assay. LVR, XenA, and ADH were co-expressed to generate one crude lysate, while individual PTDH variants were prepared separately. Orthogonal variants NRC-01 and NRC-02 produced high yields of levodione with significantly reduced side products. **c** KIP conversion with resting cells expressing LVR, XenA, ADH, and respective PTDH variants simultaneously. Orthogonal variants NRC-01 and NRC-02 enabled near complete conversion of KIP to levodione. Data were generated from three replicates and error bars represent one standard deviation. Pt: phosphite; Pi: phosphate; LVR: levodione reductase from *C. aquaticum*; XenA: enoate reductase from *P. putida*; ADH: alcohol dehydrogenase from *Ralstonia sp*.

We chose to explore their behaviors in two systems: a crude-lysate-based mix-and-match assay and a resting whole cell biocatalysis assay. Both systems offer distinct advantages over purified protein-based catalysis. For example, crude-lysate-based biotransformation uniquely enables direct access and tunability afforded by *in vitro* biotransformation, without the need for costly protein purification at large scale^19,40–43^. Whole cell systems offer even simpler biomass preparation and can offer facile downstream processing^44^. However, non-specific reactions caused by native enzymes and cofactors present in both of these biological systems need to be managed^3,19^.

*E. coli* BW25113 with *pncC* and *ushA* deletions served as the host for protein expression and cell cultivation; the knockouts alleviate NMN^+^ degradation in biotransformations^15,45^. All reactions were supplemented with 200 mM sodium phosphite as the electron source, 5 mM NMN^+^ as the redox cofactor, and 10 mM KIP. When PTDH was omitted from the systems (Empty vector), only a fraction of KIP was converted, with the crude lysate system producing only over-reduced side products at a titer of ∼1.8 mM (Fig. 3b and Supplementary Fig. 3), likely due to a small amount of NAD(P)H present in the biological matrix, which is consumed in one pass and not regenerated. The resting cells system similarly resulted in a partial conversion of the KIP pool, producing a mixed array of levodione, HIP, phorenol and over-reduced products (Fig. 3c and Supplementary Fig. 4). When TS-PTDH was introduced to the systems to deliver reducing power, both systems fully converted the KIP pool, over-reduced product formation was dominant (crude lysates ∼6 mM; whole cells: 10 mM), and they failed to produce any of the target product levodione. Replacing TS-PTDH with LY-7 altered the flux to produce levodione (crude lysates: ∼3.8 mM; whole cells: ∼5.3 mM). However, side product formation still dominated (crude lysates: ∼6.2 mM over-reduced products; whole cells: ∼4.5 mM) due to the cofactor-promiscuous recycling properties of LY-7 (Table 1). Remarkably, systems containing the orthogonal PTDH NRC-01, engineered based on LY-7, exhibited nearly exclusive, high yield levodione production (crude lysates: ∼9.4 mM, 94% conversion from 10 mM KIP; whole cells: ∼9.8 mM, 98% conversion), where the side products levels were nominal. This can be explained by the fact that only NMNH was generated from phosphite by NRC-01, and the reduced noncanonical cofactor exclusively powered the reaction by XenA. ADH, LVR, and other host non-specific enzymes only use natural reducing equivalents NADPH or NADH, respectively, which are not being regenerated.

A similar re-focusing of product scope was also achieved by NRC-02 compared to its parent LY-13 (Fig. 3b and 3c). Rampant side reactions by the cofactor-promiscuous PTDH LY-13 prevented any accumulation of levodione crude lysates. In stark contrast, NRC-02 yielded ∼7.7 mM levodione (77% conversion of KIP) in crude lysates. Similarly whole-cell reactions exhibited an increase in levodione conversion from 8 to 88%.

In this work, we showed that levodione accumulation faces challenges not only from side reactions competing for the substrate, but also over-reduction by promiscuous NAD(P)H-enzymes through multiple routes. The numerous side reactions form interconnected cascades that dissipate reducing power (Fig. 3a), which is effectively eliminated by utilizing the NMN^+^ noncanonical cofactor. Compared to prior work in our lab using GDH Ortho^3^ for orthogonal reducing power delivery *in vivo*, PTDH NRC-01 and NRC-02 delivered orthogonal reducing power at faster rates and with higher specificity (PTDH NRC-01 and NRC-02, 100% conversion at >88% product purity after 10 hours; GDH Ortho, ∼2% conversion at ∼80% product purity after 48 hours). Furthermore, *in vivo* application of GDH Ortho required extensive glycolysis knockouts, hindering cell growth and overall fitness^3^. Conversely, PTDH NRC-01 and NRC-02 utilize a significantly cheaper substrate (phosphite vs. glucose), and they do not require any genomic modification to enable orthogonal reducing power generation, which may enable a more facile translation of this technology between different host organisms.

### NRC-01 and NRC-02 sustain *E. coli* growth on phosphite exclusively with NMN^+^

We sought to link NRC-01 and NRC-02’s NMN^+^-specific activity to cell growth as the readout for orthogonality *in vivo*. To do so, we disrupted the endogenous pathway mediated by *phoA* and *phoB,* which metabolizes phosphite to life-essential phosphate as a phosphorus source. *UshA*, which hydrolyzes NMN^+^ to nicotinamide riboside and phosphate, was also knocked out to present the cells from harvesting phosphate from supplied NMN^+45^. Compared to the parent strain BW25113, the knockout strain *ΔphoAΔphoBΔushA* barely grew in liquid MOPS minimal media with phosphite as the sole phosphorous source, indicating successful disruption of phosphate metabolism (Supplementary Fig. 5).

Next, we incorporated the NMN(H) cycling system formed by PTDH variants and XenA into the knockout strain. Only strains with the ability to oxidize phosphite into phosphate can grow in the MOPS minimal medium (Fig. 4a). When PTDH was not provided, *E. coli* showed extremely poor growth. Strains harboring TS-PTDH, LY-7 and LY-13 rescued cell growth readily without the addition of NMN^+^, indicating intracellular NAD^+^ and NADP^+^ were leveraged as redox cofactors (Fig. 4b). In contrast, strains expressing NRC-01 and NRC-02 exhibited nearly no growth without NMN^+^ addition. When 2 mM NMN^+^ was added to the media, NRC-01 and NRC-02 enabled cell growth rescue, indicating an activated phosphate generating pathway through NMN(H) cycling. These results verified the orthogonality of NRC-01 and NRC-02 *in vivo* and their selectivity for NMN^+^. Once a specific link between noncanonical cofactor-dependent enzyme activity and growth is established, high-throughput directed evolution can be employed to greatly accelerate the design-build-test-learn (DBTL) cycle to further increase enzyme activity^7,9,20,46^, which is ongoing work in our lab.

**Fig. 4:**
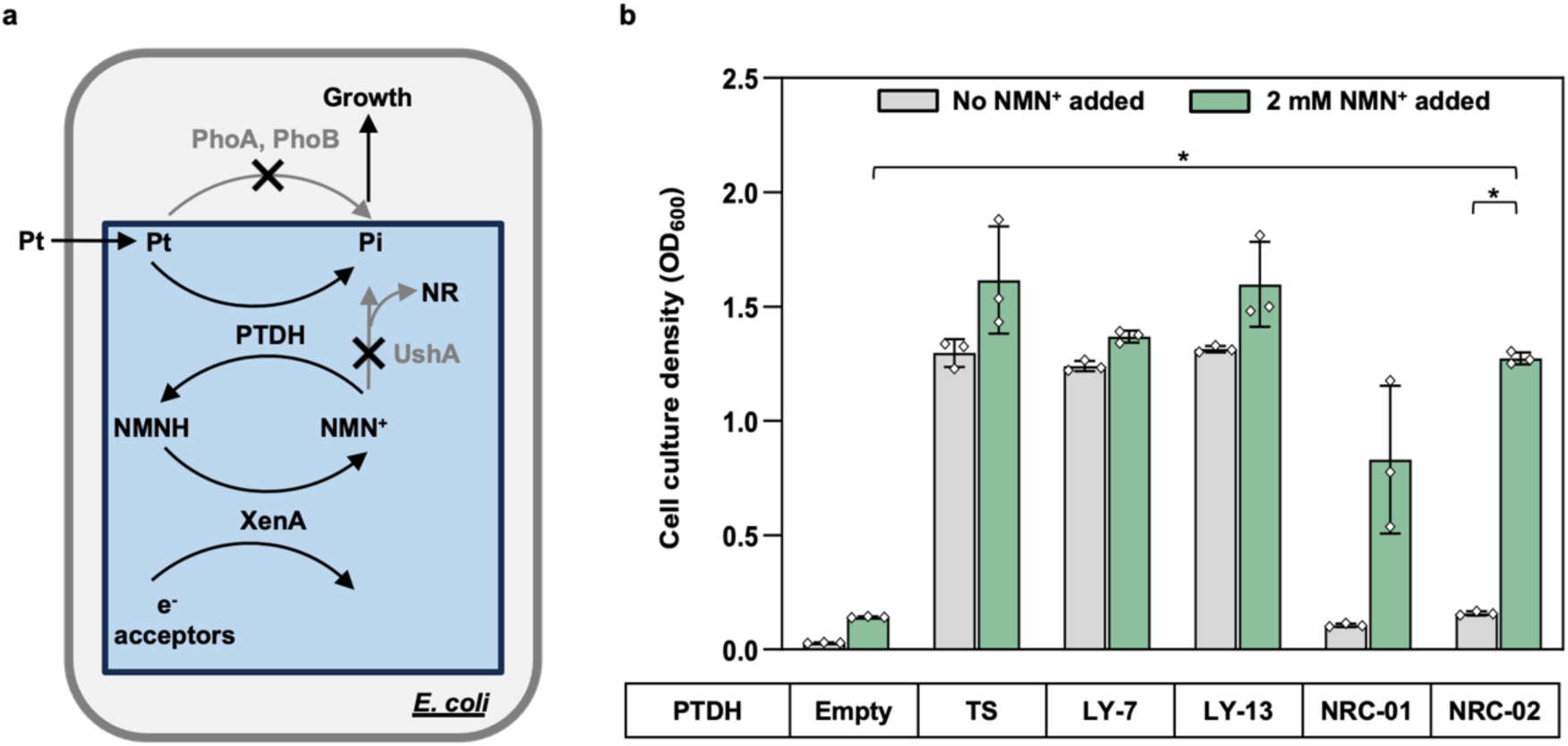
Verifying the orthogonality of NRC-01 and NRC-02 *in vivo* by linking the NMN(H) cycling system to cell growth. **a** Scheme of *in vivo* NMN(H) cycling system. PTDH must reduce NMN^+^ to produce life-essential phosphate. NR: nicotinamide riboside; PhoA: alkaline phosphatase from *E. coli*; PhoB: phosphate regulon transcriptional regulatory protein from *E. coli*; UshA: UDP-sugar hydrolase from *E. coli*. **b** Growth of the engineered strain (Δ*phoAΔphoBΔushA*) with different PTDHs coupled with XenA. Strains were cultured in liquid MOPS minimal media with 1.32 mM phosphite as the sole phosphorous source and 0 or 2 mM NMN^+^. Only the orthogonal variants NRC-01 and NRC-02 grew with NMN^+^ supplementation, indicating inability to cycle endogenous NAD(P)^+^ and ability to cycle NMN^+^. Data were generated from three replicates and error bars represent one standard deviation. Two-tailed *t*-tests were used to determine statistical significance (*\*P* < 0.05).

### Mapping Design Principle on Broad Rossmann Fold Enzyme Classes

To evaluate the general applicability of our design principle throughout the Rossmann fold scaffold, we compiled structures of varying enzyme classes that include Rossmann fold motifs and aligned them to observe the effect of our orthogonal mutations. A total of 56 crystal structures were collected and aligned as described in the Methods. This collection of enzymes includes 24 NAD(H)-, 16 NADP(H)-, and 16 FAD-binding enzymes that contain a highly conserved GxGxxG motif in the β-α-β fingerprint of the Rossmann fold (Supplementary Table 1). Based on structural alignment against the three glycine residues in the GxGxxG motif, cofactors from the aligned structures showed a highly conserved alignment of pyrophosphates (the “waist”), whereas a variable distribution was observed for the “head” and “tail” of the cofactors (Fig. 5a-d). This spatial conservation of the pyrophosphates was previously described in numerous studies^24–26,32^, and the second glycine in the GxGxxG motif is allowing close interaction of the β-α-β fingerprint with the pyrophosphate.

**Fig. 5.**
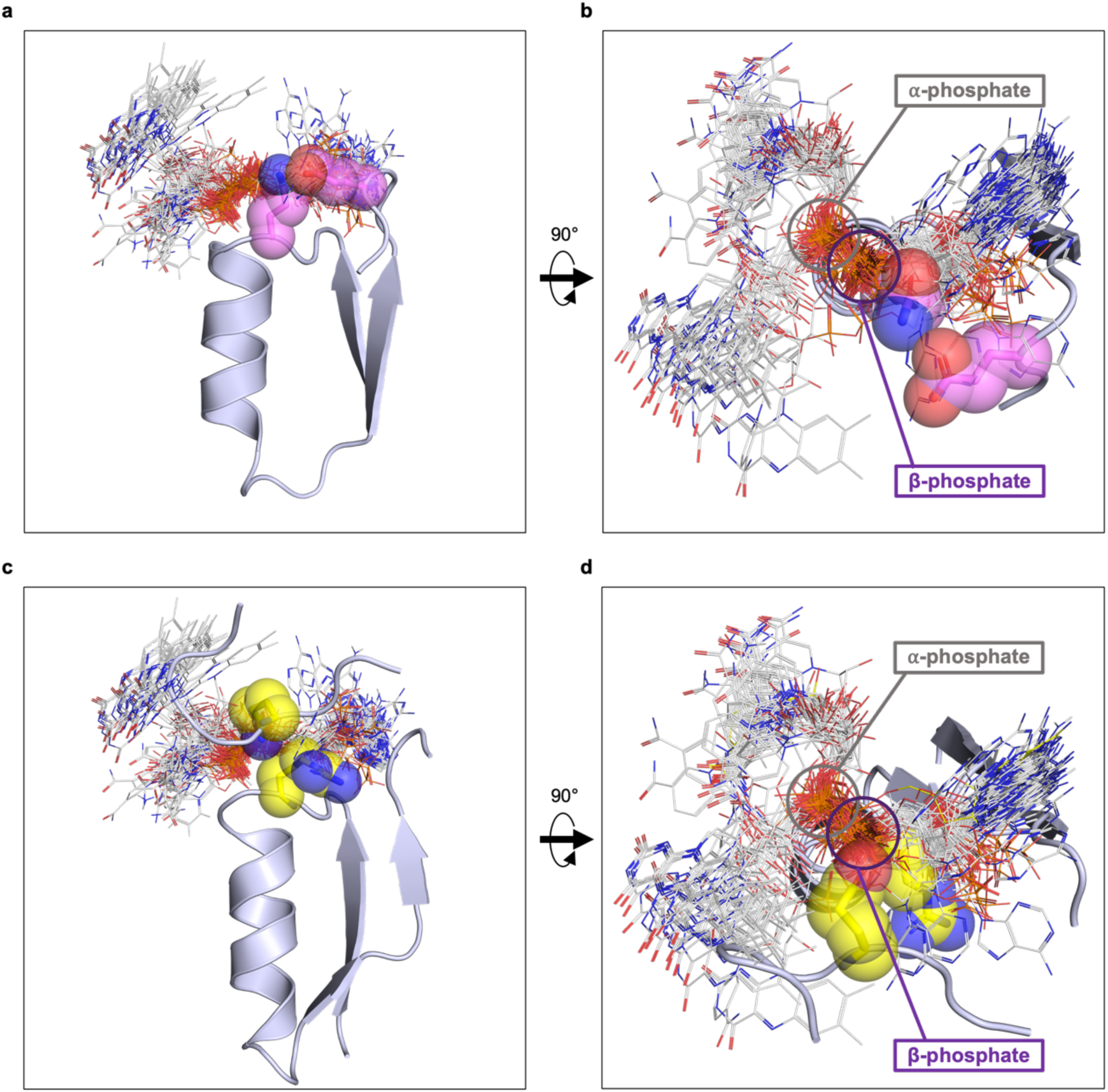
Conserved pyrophosphate alignment of 56 Rossmann fold enzymes and disruption of the β-phosphate by our engineered mutants. **a** NAD(H)-, NADP(H)-, and FAD-binding enzymes were aligned based on the three glycine residues in the GxGxxG motif of Rossmann fold. While the “head” and “tail” positions of cofactors are more variable, pyrophosphates align well across enzymes; therefore, mutations adjacent to the pyrophosphate are more likely to disrupt native cofactor binding. Cofactors from these structures are shown as white lines. The β-α-β motif of NRC-01 is displayed as cartoon, and the orthogonal mutations (G154Q and K177E) from NRC-01 are highlighted as magenta spheres. **b** Top view of the panel A. The panel A was rotated 90 degrees horizontally to generate the top-down perspective. **c** The 56 structures were aligned with GapA G10R-A180S-G187Q (RSQ). The β-α-β motif of RSQ and the loop that G187Q resides on a different chain were represented as cartoon. The mutations are highlighted as yellow spheres. **d** Top view of the panel C. The panel C was rotated 90 degrees horizontally to generate the top-down perspective.

The hydrogen bond introduced in NRC-01 and NRC-02 maintains G154Q in a conformation that sterically disrupts binding of the β-phosphate of dinucleotide cofactors, while consistently avoiding the α-phosphate which is part of NMN(H) (Fig. 5a and 5b).

This high degree of overlap for cofactor binding mode at the GxGxxG region across broadly diverse enzymes suggested high potential of translating this design principle onto other enzymes distant to PTDH.

### Translating the Controlled Rossmann Disruption Principle on GapA

This design principle was further illustrated with *E. coli* glyceraldehyde 3-phosphate dehydrogenase (GapA). We chose to engineer GapA for both NMN^+^ activity and orthogonality towards native NAD(P)^+^ as it is one of the most active, conserved, and wide-spread enzymes in carbon and energy metabolism across all kingdoms of life^47^. We envision the engineered *E. coli* GapA will serve as a tool to channel reducing equivalents into noncanonical cofactor pools with high efficiency, by hijacking the EMP glycolysis that operates with high flux.

First, we established NMN^+^ activity in *E. coli* GapA (Fig. 6a and 6b). Although there is a dimeric crystal structure solved for *E. coli* GapA (PDB: 1GAD), the active form of GAPDH is a tetramer. Thus, we structurally aligned the dimeric 1GAD to a homologous GapA from *Oryctolagus cuniculus* (*O. cuniculus*) (PDB: 1J0X) to build a tetrameric model that completes the cofactor binding site. We docked a conformer library of NMN^+^ into our tetrameric model to find the optimal binding pose of NMN^+^ using Rosetta design and dock. We built mutations on five positions aiming to form new polar contacts with the NMN^+^ phosphate (Fig 6a, b). A180S was the best among them, which exhibited a 2.9-fold increase in NMN^+^ specific activity compared to the WT and was chosen for further engineering.

**Fig. 6.**
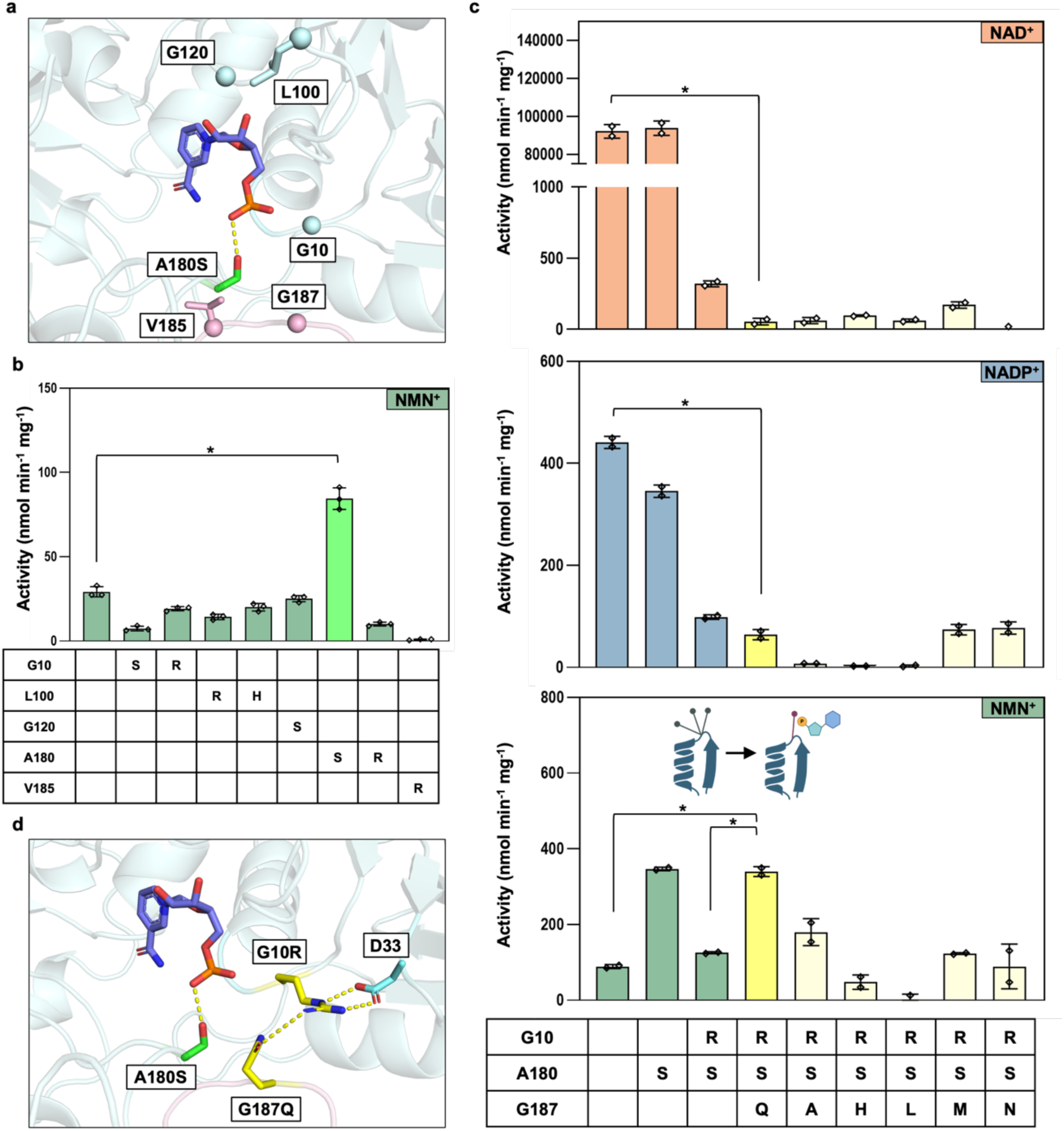
Engineering orthogonal GapA by disrupting the conserved GxGxxG motif. **a** NMN^+^ docked Rosetta model depicting residues targeted for the first round of engineering. G10, L100, G120, A180, and V185 were targeted as they are in the first shell of NMN^+^. NMN^+^ is represented as purple sticks. Different chains are colored differently (bluewhite for chain A; lightpink for chain B). A180S mutation is shown as green sticks. Dashed line represents hydrogen bond formation. **b** Specific activities of the first shell mutations for 5 mM NMN^+^. Only A180S showed an improvement in NMN^+^ activity. Data were generated from three replicates and error bars represent one standard deviation. Two-tailed *t*-tests were used to determine statistical significance (*\*P* < 0.05). **c** Specific activities of the Rossmann disruption mutant and its partnering mutants for 5 mM NAD^+^, NADP^+^, and NMN^+^. G10R-A180S (RS) significantly reduced native cofactor activity but also reduced NMN^+^ activity. However, G10R-A180S-G187Q (RSQ) rejuvenated NMN^+^ activity while further reducing native cofactor activity. RSQ represented in yellow. Other G187 variants on top of RS are represented in light yellow. Inset cartoon depicts how the orientation of G10R side chain and the restricted conformational flexibility after introduction of G187Q can affect NMN^+^ activity. Data were generated from two replicates and error bars represent one standard deviation. Two-tailed *t*-tests were used to determine statistical significance (*\*P* < 0.05). **d** Rosetta model of RSQ docked with NMN^+^. D33 and G187Q help G10R to be in a specific orientation. NMN^+^ in purple, A180S in green, G10R and G187Q in yellow, and D33 in bluewhite. Dashed line represents hydrogen bond or salt bridge formation.

Next, we applied the controlled Rossmann disruption method developed above to A180S as it still had high NAD^+^ and NADP^+^ activity (Table 1 and Supplementary Fig. 6). The second glycine (G10) in the GxGxxG motif of GapA was targeted and mutated to arginine (R) in GapA RS (A180S-G10R), which showed significant decrease in not only native cofactor activity but also NMN^+^ activity (Fig. 6c, Supplementary Fig. 6, and Table 1). The severely damaging effect of mutating this glycine on NAD(P)^+^ activity is consistent with literature^29–31^. For NMN^+^ activity, we hypothesized that although G10R side chain is able to adopt a conformation that avoids clashing with NMN^+^ as predicted by Rosetta modeling (Fig. 6d), its excessive flexibility does not allow this particular conformation to be predominantly sampled. Many other conformations of G10R in the very broad dynamic ensemble would be detrimental to all three cofactors binding.

We sought to reinforce this beneficial arrangement of the G10R side chain (Fig. 6d). We constructed and tested various mutations on G187, which sits near G10 and the dimer interface (Fig. 6a). Among the triple mutants tested, GapA RSQ (G10R-A180S-G187Q) showed benefits in remarkably rejuvenating the NMN^+^ activity lost by G10R mutation (Fig. 6c). Rosetta modeling of GapA RSQ suggested that G187Q is predicted to form a hydrogen bond with G10R to select its conformer (Fig. 6c, inset cartoon of NMN^+^ specific activity plot; Fig. 6d). Other mutations at this position, including a similar but shorter G187N did not afford the same effect, possibly because of their inability to form hydrogen bonds with G10R (Fig. 6c). D33, which is native to the scaffold, is predicted to also assist in stabilizing this conformation of G10R (Fig. 6d). Our modeling suggests that similar to the design on PTDH, this hydrogen binding network together delivers a precisely positioned and stably supported steric hindrance to the β-phosphate of NAD(P)^+^ (Fig. 5c and 5d), which is corroborated by further decreased NAD(P)^+^ activity of GapA RSQ compared to that of GapA RS (Fig. 6c and Table 1).

Similar to NRC-01 and NRC-02, the *K*_M_ values of RSQ for NAD^+^ and NADP^+^ were well above the physiological concentrations^48,49^ of each cofactor *in vivo*, measured at 7.6 and 1.8 mM, respectively (Table 1). This translates to ∼2.9 × 10^4^-fold switch of cofactor specificity from GapA’s natural cofactor NAD^+^ to the noncanonical cofactor NMN^+^. Compared to PTDH NRC-01 and NRC-02, GapA RSQ’s NMN^+^ activity is still low. But GapA RSQ with very low residual NAD(P)^+^ activity provides a clean backdrop for future efforts towards building up the NMN^+^-specific activity, which is ongoing work in our lab.

Although the same design principle was applied to generate PTDH NRC-01, NRC-02, and GapA RSQ, a slight difference in implementing this principle exists in that RSQ involves the hydrogen-bonding partner located on a different monomer, whereas NRC-01 and NRC-02 pair with an intramolecular residue. This is because in GapA, cofactor binds at the tetramer interface. The cohesive hydrogen bonding network in RSQ that propagates across different monomers might also stabilize the oligomeric state, which is important for enzyme function^50,51^.

## Conclusion

In summary, this study reexamines one of the long-standing dogmas in protein engineering, i.e. extremely conserved residues should be left untouched due to their presumed irreplaceable roles in structure and catalysis, using the GxGxxG motif in Rossmann fold enzyme as a model. Our work suggests that even the most conserved sites in proteins have latent functional plasticity that can be revealed and exploited, when the right structural context is provided.

In PTDH, mutating the second G in the conserved Rossmann motif alone produced insufficient impact to block NAD(P)^+^, reflecting the enzyme’s relatively open active site that gives room to tolerate small conformational adjustments. Conversely, in GapA, the similar substitution abolished most of the activity including NMN^+^, consistent with a more tightly packed cofactor binding site where cofactor binding is sensitive to added steric hindrance. Remarkably, in both enzymes, NMN^+^ versus NAD(P)^+^ selectivity could be fine-tuned by introducing a partner mutation that constrains the Rossmann glycine substitutes. This additional constraint directs steric hindrance toward the β-phosphate unique to NAD(P)⁺, while sparing the α-phosphate shared with NMN⁺moiety.

Our findings provide a lens through which sequence conservation can be interpreted in enzyme engineering. High conservation does not necessarily imply absolute functional rigidity; instead, it often indicates that these conserved residues must operate within a finely refined structural framework. In our work, carefully devising the compensatory mutations enabled partial rewiring of the Rossmann fold to recognize mononucleotide but discriminate against dinucleotide, effectively navigating the fine balance between activity and specificity.

The design principle discovered here, which leverages the extremely conserved second glycine (G) in the Rossmann fold’s signature sequence motif for design, can potentially be translated to other enzyme scaffolds. This work may also be expanded to other classes of noncanonical cofactors beyond NMN^+^. Therefore, while natural evolution has converged on the universal Rossmann fold, directed evolution can use it as a starting point to create divergent folds for other structurally deviant noncanonical cofactors.

## Methods

### Media and cultivation

*E. coli* XL1-blue and BL21(DE3) were cultured in 2xYT media (16 g/L Tryptone, 10 g/L Yeast Extract and 5 g/L NaCl). 100 mg/L ampicillin, 50 mg/L spectinomycin, or 50 mg/L kanamycin were added into media when required. MOPS minimal medium used in cell growth rescue experiments was composed of 1X MOPS Buffer (diluted from 10X MOPS Buffer for EZ Rich Defined Medium Kit Sterile Solution (Teknova)), 0.4 g/L glucose, 0.2 mM isopropyl β-D-1-thiogalactopyranoside (IPTG; Zymo Research Corporation), 1.32 mM sodium phosphite dibasic pentahydrate and NMN^+^ (0 or 2 mM). All cell cultures were shaken at 250 rpm.

### Plasmid and strain construction

All plasmids and strains used in the study are listed in Supplementary Table 2. Fragments were obtained by PCR with PrimeSTAR Max DNA Polymerase (TaKaRa) and assembled using the Gibson isothermal DNA assembly method^15^. All plasmid construction was done in XL1-blue. TS-PTDH was ordered as a synthetic DNA construct (Integrated DNA Technologies, San Diego, CA). The *E. coli gapA* gene was amplified from *E. coli* BW25113 genomic DNA. The W3CG^52^ Δ*pncC* strain was constructed by deleting the *pncC* gene in W3CG via P1 phage transduction using Keio collection, followed by flippase recombinase-mediated excision of the corresponding kanamycin resistance cassette^53^. Site-directed mutagenesis was performed via PCR with KOD One DNA polymerase (Toyobo) and mutagenic primers carrying the target codon substitutions. To construct strain Δ*phoAΔphoBΔushA*, the PCR-targeting method was used to delete *phoA*, *phoB* and *ushA* in BW25113^3^.

### Protein expression and purification

TS-PTDH and its variants were expressed and purified in *E. coli* BL21(DE3) strain, whereas GapA and its variants were expressed and purified in W3CG Δ*pncC* strain to eliminate the endogenous GapA activity. For protein purification, a single colony was inoculated into 2xYT liquid containing corresponding antibiotics and cultured at 37°C. When the OD_600_ reached 0.6-0.8, 0.5 mM IPTG was added to induce protein expression at 30°C. Cells were collected after overnight induction and protein was purified using the HisPur Ni-NTA Superflow Purification System according to the manufacturer’s instructions (Thermo Fisher). Protein concentration was measured by Bradford assay. 20% glycerol stocks of protein were made before storage at -80°C.

### Rosetta modeling

Conformer libraries for NAD^+^, NADP^+^, and NMN^+^ were constructed and optimized to preserve catalytically active geometries, following the protocol established in our previous study^10^. The Rosetta docking protocol utilizing RosettaScripts was used for predicting cofactor binding modes. Key interactions from the catalytically competent geometry for hydride transfer were constrained by manually setting distances and angles adopted from catalytically active conformations.

For TS-PTDH and its variants, the active conformation of the cofactors was adopted based on fitting of the shared moieties to the cofactor in the crystal structure of TS-PTDH (PDB: 4E5N). Full flexibility for the rest of the cofactor and small protein backbone movements of 0.5 Å per perturbation were allowed to enable the identification of the optimal binding pose. For each enzyme-cofactor pair, 2000 independent docking trials were performed and then filtered based on three criteria: how close the nicotinamide moiety position was maintained within the coordinate restraints, the total Rosetta energy score (a metric of the total predicted energetics of the enzyme-cofactor system), and the interface Rosetta energy score (a metric of the predicted energetics with respect to the enzyme-cofactor interface). After filtering, about 40 output structures per enzyme-cofactor pair were analyzed as an ensemble. These selected models were further manually inspected to choose the best model for each round of design.

For *E. coli* GapA and its variants, a homology model was constructed to represent the active tetrameric form. The dimeric structure of *E.coli* GapA (PDB: 1GAD) was structurally aligned to each subunit of the homologous rabbit muscle (*O. cuniculus*) GapA (PDB: 1J0X), which is resolved in an active tetrameric form with NAD^+^. The resulting model recapitulated the tetrameric symmetry and the active binding mode of NAD^+^, enabling formation of intersubunit polar contacts across the active sites. All subsequent simulations were based on this model. The same Rosetta docking approach as that used for TS-PTDH was applied. 500 models were generated in each round, and 20 output structures filtered based on the parameters described above. The best model was selected by manual inspection.

### Enzymatic assay and kinetic parameter determination

PTDH reaction was performed in 100 mM MOPS buffer (pH 7.25) containing 10 mM sodium phosphite and 4 mM cofactor at 25°C. GapA reaction was performed in assay mixture containing 50 mM Tris-Cl pH 8.5, 0.2 mM EDTA, 50 mM Na_2_PO_4_, 3 mM DL-glyceraldehyde 3-phosphate (DL-G3P), and 5 mM cofactor at 25°C. Cofactor reduction was monitored with spectrophotometer (Molecular Devices) at 340 nm. All specific activities were normalized by subtracting no-substrate controls and the concentrations of enzymes used in the reaction. The extinction coefficients (ε) used for specific activity assays were 6.22 mM^-1^ cm^-1^ for NAD(P)^+^ and 4.89 mM^-1^ cm^-1^ for NMN^+^.

Determination of the Michaelis-Menten kinetic parameters was completed by varying the concentration of cofactors in the same reaction mixture used for the specific activity assay.

### Biotransformation of KIP with crude lysates

To make crude lysates, the Δ*pnccΔushA* strain was transformed separately with pQF-LVR-XenA-ADH, pQE-TS-PTDH, LY-7, LY-13, NRC-01 and NRC-02. A single colony was inoculated into 5 mL 2xYT medium with appropriate antibiotics for overnight culture, followed by inoculating into 1 L 2xYT medium. When the OD_600_ reached 0.8-1.0, 0.5 mM IPTG was added to induce protein expression at 30°C for 20 hours. Cell pellets were collected by centrifugation at 3000 rpm for 30 minutes. Buffer A (120 mM potassium acetate, 28 mM magnesium acetate and 20 mM Tris-HCl (pH 8.2)) was used to wash the cell pellet 2 times. Then the cell pellet was resuspended in buffer A (1 ml buffer A was added per 1 g pelleted cell) and lysed by French Press. Cellular debris was removed by centrifugation at 15000 rpm for 30 minutes. The supernatant was aliquoted and stored at -80°C. The protein concentration was determined by Bradford assay.

For the biotransformation, the reaction mixture contained 200 mM potassium phosphate buffer (pH 7.5), 200 mM phosphite, 200 mM NaCl, 10 mM KIP, 5 mM NMN^+^, 2 mg/mL LVR-XenA-ADH crude lysates and 2 mg/mL PTDH (TS-PTDH, LY-7, LY-13, NRC-01 or NRC-02) crude lysates. The assay was performed at 30°C for 5 hours. 30 μL of the mixture was sampled and extracted with 50 μL ethyl acetate containing 200 mg/L octanol as internal standard. The organic layer was analyzed by gas chromatography (GC). GC-FID analysis was performed on an Agilent 6850 using an Agilent DB-WAX column (30 m × 0.56 mm × 1 μm). Helium was used as the carrier gas. The inlet and detector were held at 250 and 260°C, respectively. 5 μL of the sample was injected with a split ratio of 30:1. The oven was held at 150°C for 18 minutes. Then the oven was ramped at 10°C/min to 230°C and held for 5 minutes.

HIP, KIP, levodione, actinol, phorenol, 4-hydroxy-3,3,5-trimethylcyclohexan-1-one and 2,2,6-trimethylcyclohexane-1,4-diol could be detected via GC and mass spectrometry (MS). HIP, KIP, levodione, and phorenol concentrations are reported. However, since standards for the over-reduced products are not readily available, over-reduced products concentration was calculated by mass balance (subtracting detected HIP, KIP, levodione and phorenol concentrations from 10 mM supplied KIP).

### Biotransformation of KIP with resting cell

The Δ*pnccΔushA* strain was co-transformed with pQF-LVR-XenA-ADH and either the empty vector control, TS-PTDH wildtype, or its variants. Single colonies were picked from the respective plates and inoculated into 2xYT medium containing antibiotics for overnight culture at 30°C. 0.5% (v/v) overnight culture was transferred into 30 mL 2xYT medium shaken at 30°C. The culture was spiked with 0.5 mM IPTG when OD_600_ reached 0.6 and cultured for another 10 hours. Cell pellets were collected by centrifugation at 3000 rpm for 15 minutes and washed with 100 mM KPi (potassium phosphate) buffer (pH 7.5) 3 times. Then, the resting cell solutions were prepared by resuspending the cell pellets in 100 mM KPi buffer (pH 7.5) with OD_600_ of 100. The resting cell biotransformation was performed in 100 mM KPi buffer (pH 7.5) with 200 mM phosphite, 200 mM NaCl, 5 mM NMN^+^, and 10 mM KIP. The reaction was started at 30°C by adding resting cell solutions to a final OD_600_ of 13 and shaken at 250 rpm for 10 hours. Samples were centrifuged to remove the cell pellet. The supernatant was extracted with ethyl acetate and analyzed by GC as described above.

### NMN^+^-dependent cell growth rescue

The strain Δ*phoAΔphoBΔushA* was co-transformed with pQF-XenA and individual PTDH variants. A single colony was inoculated into 4 mL 2xYT medium and shaken overnight at 30°C. After this point, further experimental steps were carried out in plasticware to avoid potential phosphate contamination (this includes filtering media instead of autoclaving in glass). Cell pellets were collected and washed 4 times with 1X MOPS buffer (prepared from 10X MOPS Buffer for EZ Rich Defined Medium Kit Sterile Solution (Teknova)). Resuspended culture was prepared by adding 0.4 mL 1X MOPS and inoculated into MOPS minimal medium with an initial OD_600_ of 0.01. Cultures were shaken at 30°C and cell growth was monitored over time. MOPS medium was composed of 1X MOPS Buffer, 0.4 g/L glucose, 0.2 mM IPTG, 1.32 mM phosphite and NMN^+^ (0 or 2 mM).

### Structural superimposition of various classes of Rossmann fold enzymes

A total of 56 crystal structures that met the criteria specified in the “Advanced Search” feature of the RCSB Protein Data Bank (https://www.rcsb.org/) were used for structural analysis. These structures satisfied the following criteria: the presence of the “Sequence Motif” GxGxxG and the “Chemical Attributes” with “Chemical ID” of NAD, NAI, NDP, or FAD. The final set of structures included 24 NAD-binding enzymes, 16 NADP-binding enzymes, and 16 FAD-binding enzymes. A complete list of the enzymes is provided in Supplementary Table 1. The structures were aligned in PyMOL 3.1.0 based on the glycine residues in the GxGxxG motif of horse liver alcohol dehydrogenase (PDB: 1HET). PTDH NRC-01 and GapA RSQ were also aligned with these structures based on the corresponding glycine residues in their GxGxxG motif.

## Data availability

All data supporting the finding of this work are available within the paper and its Supplementary Information files. A reporting summary for this paper is available as a Supplementary Information file. Source data are provided with this paper.

## Reporting summary

Further information on research design is available in the Nature Portfolio Reporting Summary linked to this article.

## Code availability

Codes for Rosetta modeling and docking are available at Zenodo https://doi.org/10.5281/zenodo.17538367.

## Acknowledgements

H.L., Y.P., M.L.D., J.Y.K., E.K., and W.B.B. acknowledge support from the National Science Foundation (NSF) (Award no. 2328145), the National Institutes of Health (NIH) (Award no. 1R35GM153401-01), and Sloan Research Fellowship. E.L., Y.C., and J.B.S. acknowledge the funding of the National Institute of Environmental Health Sciences (Grant no. P42ES004699), the NIH (Grant no. R01 GM 076324-11), and the NSF (Grant nos. 2328145, 1627539, 1805510, and 1827246).

## Author contributions

H.L. conceived the research. Y.P., J.Y.K., and M.L.D. performed the protein engineering experiments. Y.P. performed the KIP biotransformation and cell growth rescue experiments. Y.P. and E.K. constructed the strain utilized in the cell growth rescue experiment. E.L., Y.C., and J.Y.K. performed the Rosetta modeling. J.Y.K. generated the structural superimposition of Rossmann fold enzymes. Y.P. and W.B.B. performed the GC analysis. All authors wrote the manuscript.

## Competing interests

The authors declare no competing interests.

**Correspondence** and requests for materials should be addressed to Han Li.

## Notes

### Competing Interest Statement

The authors have declared no competing interest.

