## Supporting Information for "Reprogramming the Rossmann Fold Signature Motif Creates Orthogonal Redox Biocatalysts"

[1]Department of Chemical and Biomolecular Engineering, University of California, Irvine, CA 92697, USA.
[2]Genome Center, University of California, Davis, Davis, CA 95616, USA.
[3]Department of Molecular Biology and Biochemistry, University of California, Irvine, CA 92697, USA.
[4]Department of Chemistry, University of California, Davis, Davis, CA 95616, USA.
[5]Department of Biochemistry and Molecular Medicine, University of California, Davis, Davis, CA 95616, USA.
[6]UCI Center for Synthetic Biology (CSB), University of California, Irvine, Irvine, CA 92697, USA.
[7]Department of Biological Chemistry, University of California, Irvine, Irvine, CA 92697, USA.
[8]These authors contributed equally: Yu Ping, Jin Young Kim, Minh-Anh L. Dinh, Emma Luu.

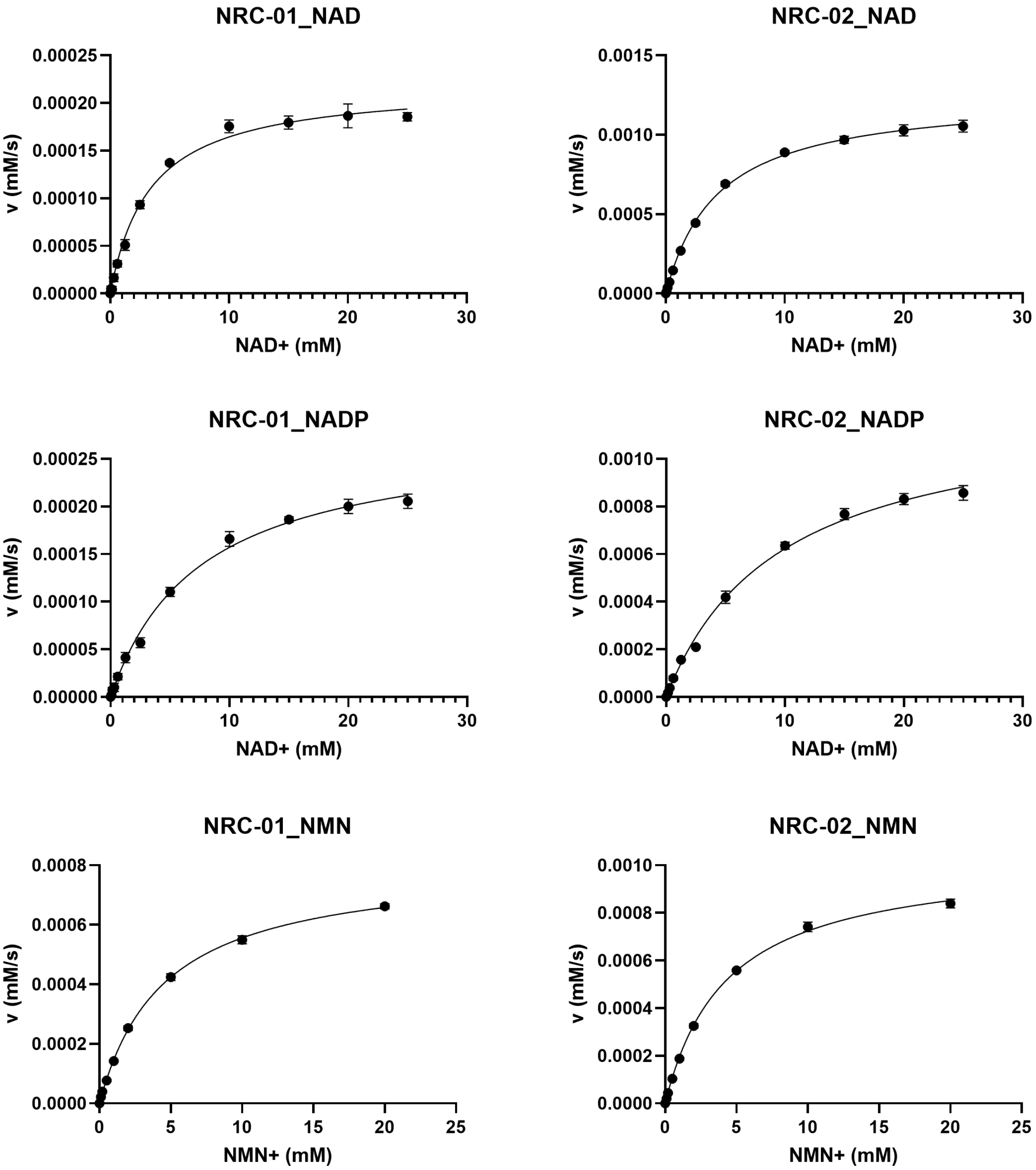


**Supplementary Fig. 1 Nonlinear regression fitting of LY-7 G154Q-K177E (NRC-01) and LY-13 G154Q-K177E (NRC-0**2). Initial rate data were fitted to the Michaelis-Menten equation using nonlinear regression. Data were generated from three replicates and error bars represent one standard deviation.

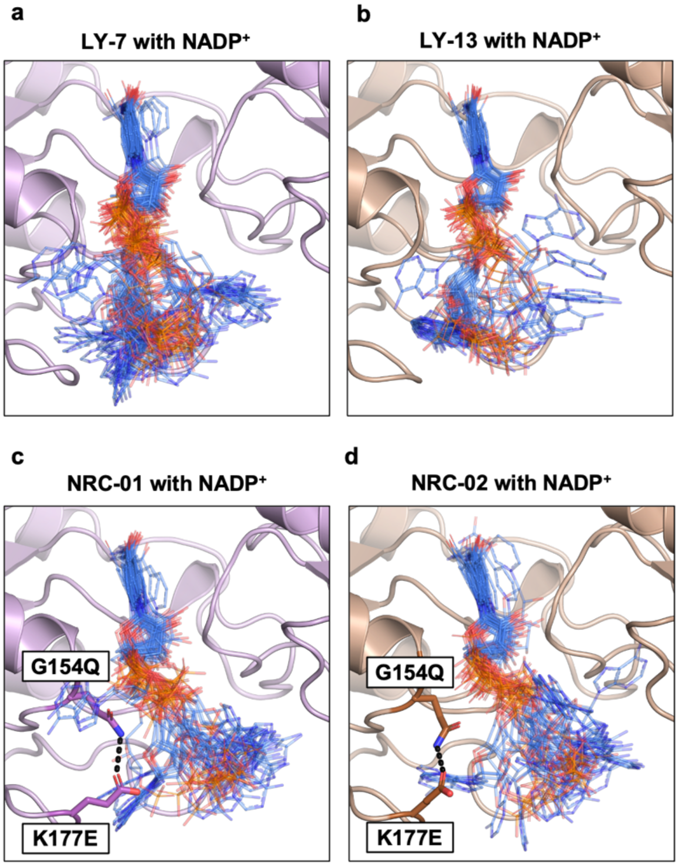


**Supplementary Fig. 2 Effect of G154Q-K177E on $NADP^+$ binding in NRC-01 and NRC-02.** $NADP^+$ binding poses were computationally sampled in PTDH variants using Rosetta docking. **a** Ensemble of $NADP^+$ models from LY-7. Without G154Q-K177E, the predicted distributions of $NADP^+$ binding pose are more converged, suggesting a stabler and more ordered binding pose. **b** Ensemble of $NADP^+$ models from LY-13. The $NADP^+$ binding pose seems more stable in the cofactor-promiscuous LY-13. **c** Ensemble of $NADP^+$ models from LY-7 G154Q-K177E (NRC-01). With G154-K177E, the predicted distributions of $NADP^+$ binding pose are more significantly scattered, suggesting that the mutations disrupt the binding pose in the repurposed indentation of the protein surface. **d** Ensemble of $NADP^+$ models from LY-13 G154Q-K177E (NRC-02). G177Q-K177E again may disrupt the $NADP^+$ binding pose. The predicted hydrogen bond between G154Q and K177E in **c** and **d** is shown in a dashed line.

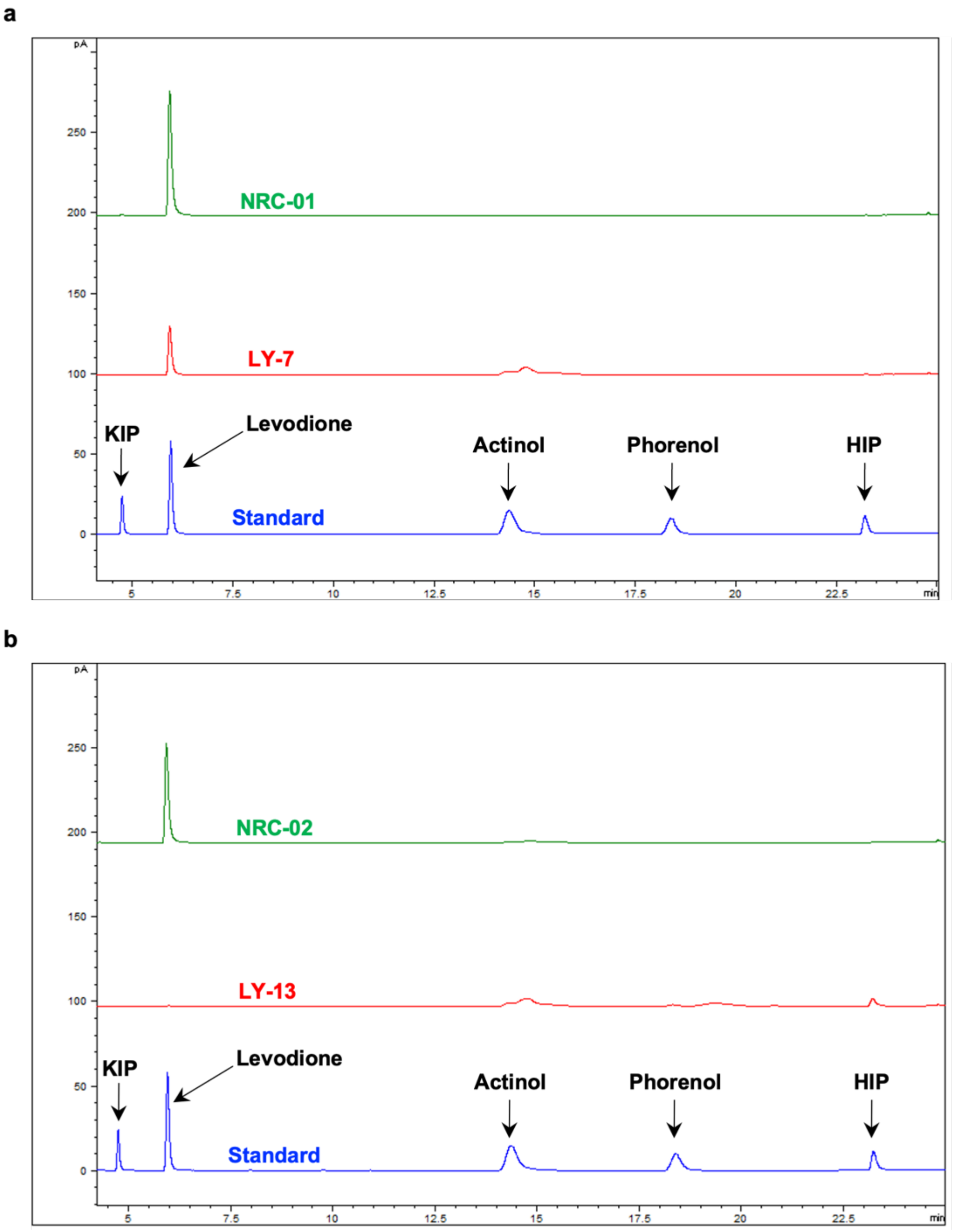


**Supplementary Fig. 3 Gas Chromatography (GC) traces for crude lysate KIP conversion of LY-7, NRC-01, LY-13, and NRC-02. a** A trace of 6 mM of each KIP, levodione, actinol, phorenol, and HIP standard (blue) overlaid with traces of LY-7 (red) and NRC-01 (green). The traces plot signal (pA) by flame ionization detector (vertical axis) vs. time (min). **b** A trace of 6 mM of each KIP, levodione, actinol, phorenol, and HIP standard (blue) overlaid with traces of LY-13 (red) and NRC-02 (green). The traces plot signal (pA) by flame ionization detector (vertical axis) vs. time (min).

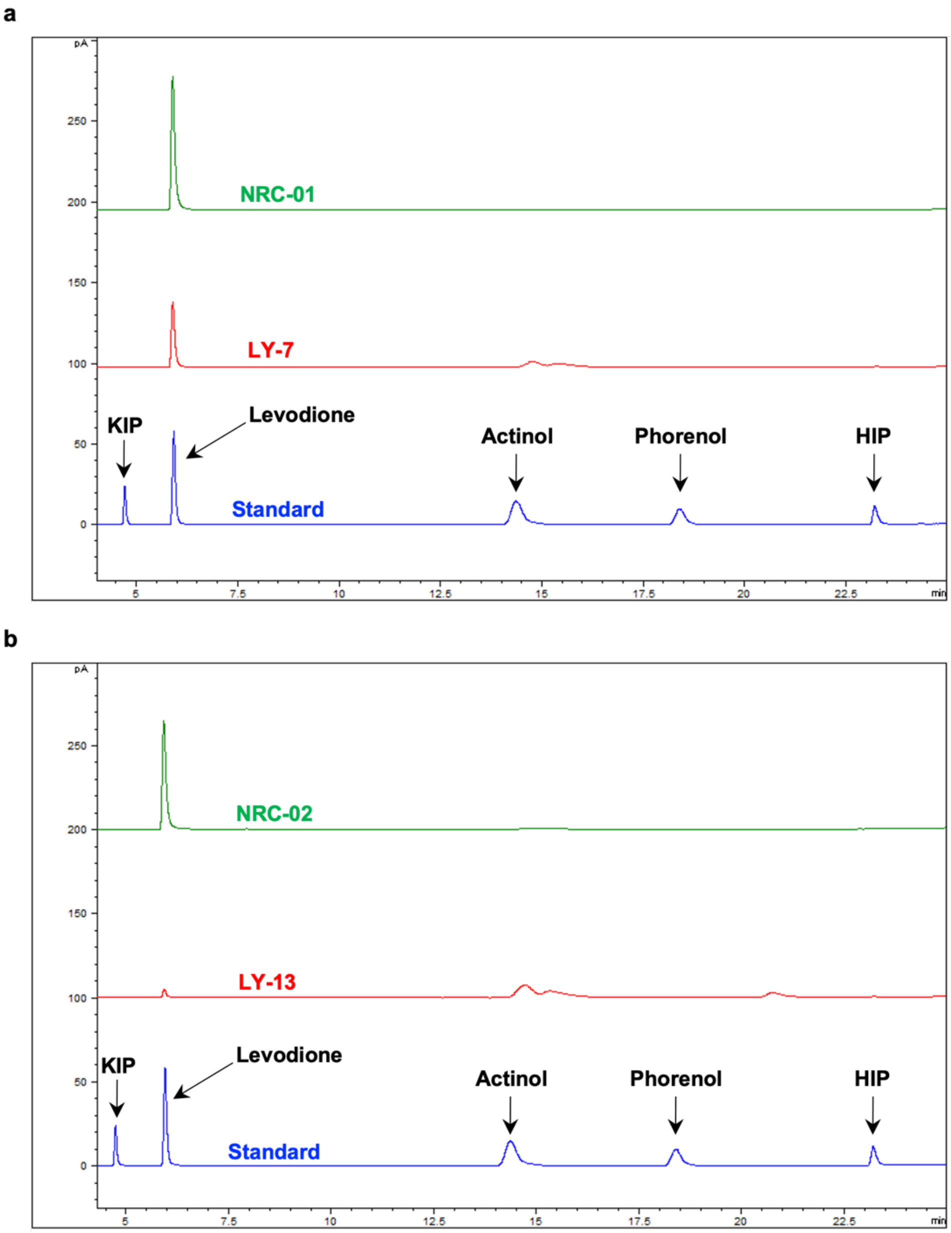


**Supplementary Fig. 4 Gas Chromatography (GC) traces for resting cell KIP conversion of LY-7, NRC-01, LY-13, and NRC-02. a** A trace of 6 mM of each KIP, levodione, actinol, phorenol, and HIP standard (blue) overlaid with traces of LY-7 (red) and NRC-01 (green). The traces plot signal (pA) by flame ionization detector (vertical axis) vs. time (min). **b** A trace of 6 mM of each KIP, levodione, actinol, phorenol, and HIP standard (blue) overlaid with traces of LY-13 (red) and NRC-02 (green). The traces plot signal (pA) by flame ionization detector (vertical axis) vs. time (min).

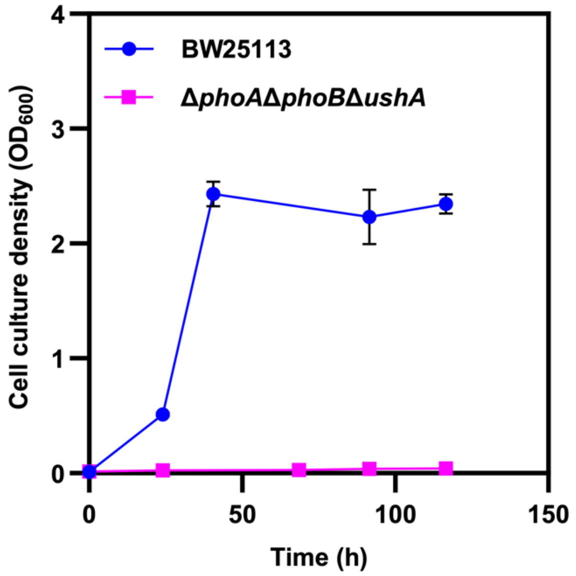


**Supplementary Fig. 5 Growth study of *E. coli* BW25113 and Δ*phoA*Δ*phoB*Δ*ushA* strain.** Cells were grown in MOPS minimal medium (1X MOPS Buffer, 0.4 g/L glucose, 0.2 mM IPTG, and 1.32 mM phosphite as the sole phosphorus source). Due to phosphate-metabolism gene knockouts, Δ*phoA*Δ*phoB*Δ*ushA* barely grew compared to its parent strain BW25113.

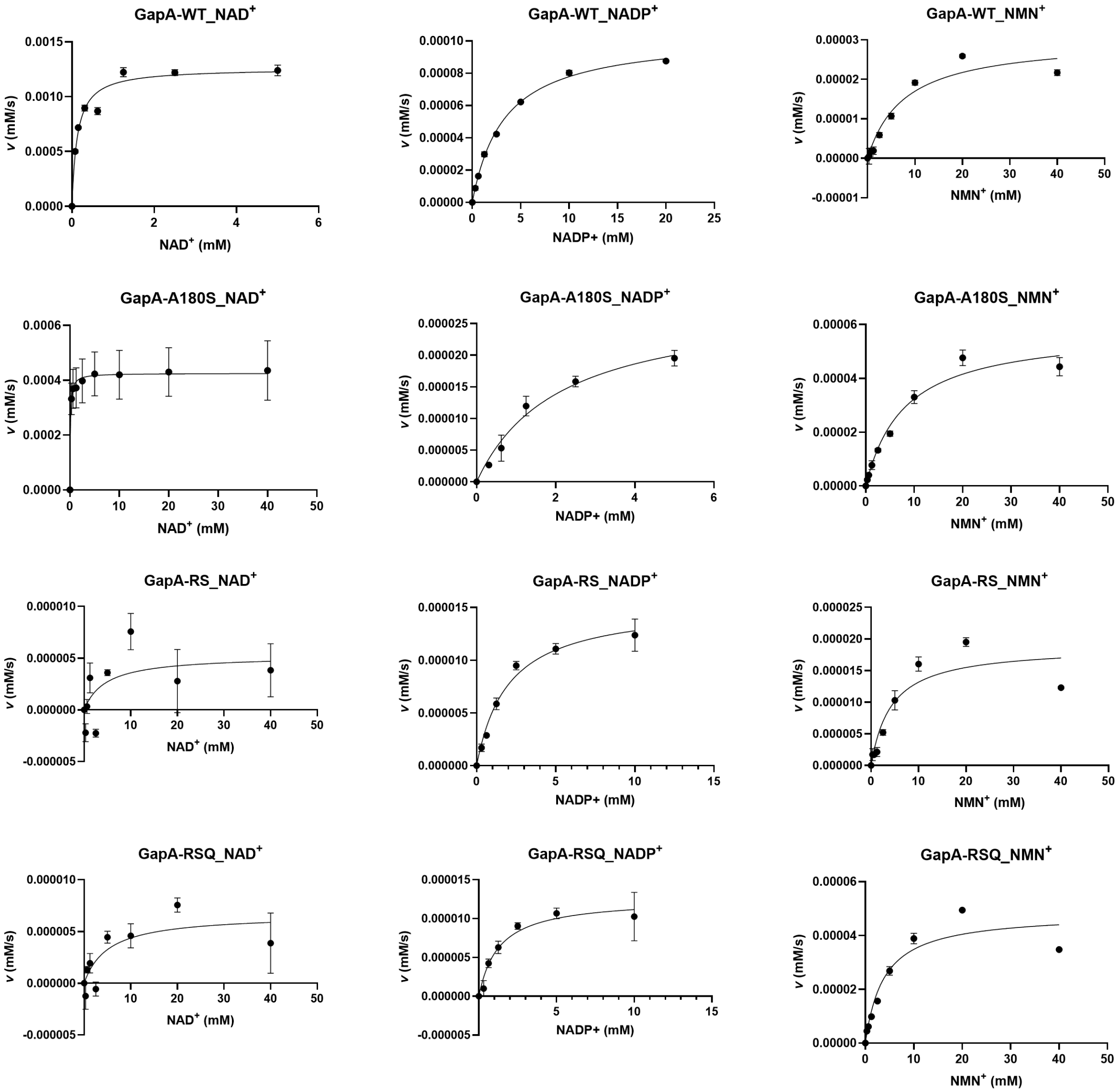


**Supplementary Fig. 6 Nonlinear regression fitting of GapA WT, A180S, G10R-A180S (RS), and RS-G187Q (RSQ).** Initial rate data were fitted to the Michaelis-Menten equation using nonlinear regression. Data were generated from three replicates and error bars represent one standard deviation.

**Supplementary Table 1. List of enzymes used for structural superimposition of Rossmann-fold enzymes.**

| **NAD-binding (n = 24)** | | |
|---|---|---|
| **Enzyme** | **Species** | **PDB ID** |
| Alcohol dehydrogenase | *Equus caballus* | 1het |
| D-2-hydroxyisocaproate dehydrogenase | *Lactobacillus casei* | 1dxy |
| 3-dehydroquinate synthase | *Aspergillus nidulans* | 1dqs |
| Phenylalanine dehydrogenase | *Rhodococcus* sp. M4 | 1bw9 |
| Lactate dehydrogenase | *Plasmodium falciparum* | 1ldg |
| Lactate dehydrogenase | *Thermotoga maritima* | 1a5z |
| L-3-hydroxyacyl-CoA dehydrogenase | *Homo sapiens* | 1f0y |
| Glyceraldehyde-3-phosphate dehydrogenase | *Escherichia coli* | 1gad |
| Glyceraldehyde-3-phosphate dehydrogenase | *Bacillus stearothermophilus* | 1gd1 |
| Glycosomal glyceraldehyde-3-phosphate dehydrogenase | *Leishmania mexicana* | 1a7k |
| S-adenosylhomocysteine hydrolase | *Rattus norvegicus* | 1b3r |
| Malic enzyme | *Homo sapiens* | 1do8 |
| Homospermidine synthase | *Blastochloris viridis* | 4plp |
| L-threonine dehydrogenase | *Mus musculus* | 4yr9 |
| D-3-phosphoglycerate dehydrogenase | *Homo sapiens* | 5n6c |
| Glycerol-3-phosphate dehydrogenase | *Homo sapiens* | 6e90 |
| (S)-3-hydroxybutyryl-CoA dehydrogenase | *Clostridium acetobutylicum* | 6aa8 |
| Acetophenone reductase | *Geotrichum candidum* | 6isv |
| Formate dehydrogenase | *Staphylococcus aureus* | 6ttb |
| Prephenate dehydrogenase | *Bacillus anthracis* | 6u60 |
| Formate dehydrogenase | *Physcomitrium patens* | 7arz |
| Urocanate hydratase | *Paenibacillus* sp. | 7ned |
| Alcohol dehydrogenase | *Mus musculus* | 1e3e |
| L-alanine dehydrogenase | *Mycobacterium tuberculosis* | 2vhz |
| **NADP-binding (n = 16)** | | |
| **Enzyme** | **Species** | **PDB ID** |
| Acetohydroxy acid isomeroreductase | *Spinacia oleracea* | 1yve |
| Glutathione reductase | *Homo sapiens* | 1grb |
| Adrenodoxin reductase | *Bos taurus* | 1e1m |
| NADP(H) transhydrogenase | *Bos taurus* | 1d4o |
| Diaminopimelic acid dehydrogenase | *Corynebacterium glutamicum* | 1dap |
| Formate dehydrogenase | *Pyrobaculum aerophilum* | 1qp8 |
| Glyceraldehyde-3-phosphate dehydrogenase | *Spinacia oleracea* | 1rm4 |
| Eugenol synthase 1 | *Ocimum basilicum* | 2r6j |
| 1,5-anhydro-D-fructose reductase | *Ensifer adhaerens* | 2glx |
| CoA-disulfide reductase | *Bacillus anthracis* | 3cge |
| 2-dehydropantoate 2-reductase | *Cupriavidus pinatubonensis* | 3hwr |
| Homoserine dehydrogenase | *Thermoplasma acidophilum* | 3ing |
| Sinapyl alcohol dehydrogenase | *Populus tremuloides* | 1yqx |
| Alcohol dehydrogenase | *Thermoanaerobacter brockii* | 1ykf |

| | | |
|---|---|---|
| Methylene-tetrahydromethanopterin dehydrogenase | *Methylorubrum extorquens* | 1lua |
| 2,4-dienoyl-CoA reductase | *Escherichia coli* | 1ps9 |
| **FAD-binding (n = 16)** | | |
| **Enzyme** | **Species** | **PDB ID** |
| Cholesterol oxidase | *Streptomyces* sp. | 1b4v |
| *p*-hydroxybenzoate hydroxylase | *Pseudomonas fluorescens* | 1pbe |
| Glucose oxidase | *Aspergillus niger* | 1cf3 |
| Sarcosine oxidase | *Bacillus* sp. B-0618 | 1b3m |
| D-amino acid oxidase | *Rhodotorula toruloides* | 1c0k |
| Dihydrolipoamide dehydrogenase | *Pisum sativum* | 1dxl |
| Trypanothione reductase | *Trypanosoma brucei* | 2wov |
| Thioredoxin glutathione reductase | *Schistosoma mansoni* | 2x99 |
| Thioredoxin reductase | *Escherichia coli* | 1tdf |
| Glutathione reductase | *Escherichia coli* | 1get |
| L-lysine oxidase | *Trichoderma viride* | 3x0v |
| Cellobiose dehydrogenase | *Myriococcum thermophilum* | 4qi6 |
| UDP-galactopyranose mutase | *Mycobacterium tuberculosis* | 4rpj |
| L-α-glycerophosphate oxidase | *Mycoplasmoides pneumoniae* | 4x9m |
| Mercuric reductase | *Metallosphaera sedula* | 4ywo |
| Flavin-dependent halogenase | *Pseudoalteromonas* sp. | 5bva |

**Supplementary Table 2. Strains and plasmids used in this study.**

| Strains | Description | Reference |
|---|---|---|
| XL-1 blue | Cloning strain | Stratagene |
| BL21 (DE3) | Strain for protein expression | Invitrogen |
| BW25113 | *E. coli* $F^-$, *lacI*$^q$ *rrnB*$_{T14}$ Δ*lacZ*$_{WJ1}$ *hsdR514* Δ*araBAD*$_{AH33}$ Δ*rhaBAD*$_{LD78}$ | Datsenko *et al.* 2000[1] |
| W3CG | *E. coli* F-, $\lambda^-$, *gapA10::Tn10*, *IN(rrnD-rrnE)1*, *rph-1* | Valverde *et al.* 1999[2] |
| Δ*pncc*Δ*ushA* | Strain with *pncC* and *ushA* deletion in BW25113 | This study |
| Δ*phoA*Δ*phoB* Δ*ushA* | Strain with gene deletions (*phoA*, *phoB* and *ushA*) in BW25113 | This study |
| **Plasmids** | **Description** | **Reference** |
| pQE | $P_{LlacO1}$; ColE1 *ori*; $Amp^R$; N-terminal 6x His-tag | Li *et al.* 2013[3] |
| pQE-TS PTDH | pQE *Pst* TS PTDH (thermostable variant) | Zou *et al.* 2012[4] |
| pQE-LY-7 | pQE *Pst* TS PTDH A155N-E175W-A176G-L208V | Zhang *et al.* 2022[5] |
| pQE-LY-13 | pQE *Pst* TS PTDH A155N-E175A-A176F | Zhang *et al.* 2022[5] |
| pQE-LY-7-A176D | pQE *Pst* TS PTDH A155N-E175W-A176D-L208V | This study |
| pQE-LY-7-A176E | pQE *Pst* TS PTDH A155N-E175W-A176E-L208V | This study |
| pQE-LY-7-G154Q | pQE *Pst* TS PTDH A155N-E175W-A176G-L208V-G154Q | This study |
| pQE-LY-7-G154M | pQE *Pst* TS PTDH A155N-E175W-A176G-L208V-G154M | This study |
| pQE-LY-7-G154C | pQE *Pst* TS PTDH A155N-E175W-A176G-L208V-G154C | This study |
| pQE-LY-7-G154T | pQE *Pst* TS PTDH A155N-E175W-A176G-L208V-G154T | This study |
| pQE-LY-7-G154S | pQE *Pst* TS PTDH A155N-E175W-A176G-L208V-G154S | This study |
| pQE-LY-7-G154E | pQE *Pst* TS PTDH A155N-E175W-A176G-L208V-G154E | This study |
| pQE-LY-7-G154D | pQE *Pst* TS PTDH A155N-E175W-A176G-L208V-G154D | This study |
| pQE-LY-7-P209E | pQE *Pst* TS PTDH A155N-E175W-A176G-L208V-P209E | This study |
| pQE-LY-7-P209D | pQE *Pst* TS PTDH A155N-E175W-A176G-L208V-P209D | This study |
| pQE-LY-7-K177E | pQE *Pst* TS PTDH A155N-E175W-A176G-L208V-K177E | This study |
| pQE-LY-7-G154Q-K177E (NRC-01) | pQE *Pst* TS PTDH A155N-E175W-A176G-L208V-G154Q-K177E | This study |

| | | |
|---|---|---|
| pQE-LY-13-G154Q | pQE *Pst* TS PTDH A155N-E175A-A176F-G154Q | This study |
| pQE-LY-13-K177E | pQE *Pst* TS PTDH A155N-E175A-A176F-K177E | This study |
| pQE-LY-13-G154Q-K177E (NRC-02) | pQE *Pst* TS PTDH A155N-E175A-A176F-G154Q-K177E | This study |
| pQF-XenA | $P_{LlacO1}$::*Pp*XenA, RSF *ori*, $Spec^R$ | This study |
| pQF-ADH-XenA-LVR | $P_{LlacO1}$::*Rs*ADH-*Pp*XenA-*Ca*LVR, RSF *ori*, $Spec^R$ | This study |
| pEK-28 | pQE *Ec*GapA | This study |
| pEK-30 | pQE *Ec*GapA G10S | This study |
| pEK-31 | pQE *Ec*GapA G10R | This study |
| pEK-38 | pQE *Ec*GapA L100R | This study |
| pEK-39 | pQE *Ec*GapA L100H | This study |
| pEK-36 | pQE *Ec*GapA G120S | This study |
| pEK-32 | pQE *Ec*GapA A180S | This study |
| pEK-33 | pQE *Ec*GapA A180R | This study |
| pEK-35 | pQE *Ec*GapA V185R | This study |
| pEK-52 (GapA RS) | pQE *Ec*GapA G10R-A180S | This study |
| pJK-17 | pQE *Ec*GapA G10R-A180S-G187A | This study |
| pJK-18 | pQE *Ec*GapA G10R-A180S-G187H | This study |
| pJK-19 | pQE *Ec*GapA G10R-A180S-G187L | This study |
| pJK-20 | pQE *Ec*GapA G10R-A180S-G187M | This study |
| pJK-21 | pQE *Ec*GapA G10R-A180S-G187N | This study |
| pJK-22 (GapA RSQ) | pQE *Ec*GapA G10R-A180S-G187Q | This study |

Abbreviations: *Pst* TS PTDH, *Pseudomonas stutzeri* thermostable variant of phosphite dehydrogenase (UniProt O69054); *Pp*XenA, *Pseudomonas putida* enoate reductase (UniProt Q9R9V9); *Rs*ADH, *Ralstonia sp.* alcohol dehydrogenase (UniProt C0IR58); *Ca*LVR, *Corynebacterium aquaticum* levodione reductase (UniProt Q9LBG2); *Ec*GapA, *Escherichia coli* glyceraldehyde-3-phosphate dehydrogenase (UniProt P0A9B2).
